# Beyond guilty by association at scale: searching for causal variants on the basis of genome-wide summary statistics

**DOI:** 10.1101/2024.02.28.582621

**Authors:** Zihuai He, Benjamin Chu, James Yang, Jiaqi Gu, Zhaomeng Chen, Linxi Liu, Tim Morrison, Michael E. Belloy, Xinran Qi, Nima Hejazi, Maya Mathur, Yann Le Guen, Hua Tang, Trevor Hastie, Iuliana Ionita-laza, Emmanuel Candès, Chiara Sabatti

## Abstract

Understanding the causal genetic architecture of complex phenotypes will fuel future research into disease mechanisms and potential therapies. Here, we illustrate the power of a novel framework: it detects, starting from summary statistics, and across the entire genome, sets of variants that carry non-redundant information on the phenotypes and are therefore more likely to be causal in a biological sense. The approach, implemented in open-source software, is also computationally efficient, requiring less than 15 minutes on a single CPU to perform genome-wide analysis. Through extensive genome-wide simulation studies, we show that the method can substantially outperform existing methods in false discovery rate control, statistical power and various fine-mapping criteria. In applications to a meta-analysis of ten large-scale genetic studies of Alzheimer’s disease (AD), we identified 82 loci associated with AD, including 37 additional loci missed by conventional GWAS pipeline. Massively parallel reporter assays and CRISPR-Cas9 experiments have confirmed the functionality of the putative causal variants our method points to. Finally, we retrospectively analyzed summary statistics from 67 large-scale GWAS for a variety of phenotypes. Results reveal the method’s capacity to robustly discover additional loci for polygenic traits and pinpoint potential causal variants underpinning each locus beyond conventional GWAS pipeline, contributing to a deeper understanding of complex genetic architectures in post-GWAS analyses.

## Introduction

Uncovering the precise causal genetic determinants of complex traits is integral to advancing the understanding of disease etiologies and the development of targeted therapies. An increasing body of evidence underscores the efficacy of therapeutic interventions grounded in genetic insights.^1,2^ Notably, approximately two-thirds of the FDA-approved new drugs in 2021 are based on genetic loci linked to their respective therapeutic indications or related phenotypes.^3^ Genome-wide association studies (GWAS) serve as one of the most popular methodologies for identifying genetic variants correlated with disease phenotypes. Nonetheless, the genetic variants identified through GWAS generally account for a limited proportion of heritability and are mostly proxies for causal variants, which may impede their applicability in functional genomics and the informed selection of drug targets and indications.^4,5^

Until recently, most GWAS have predominantly focused on marginal association models with family-wise error rate (FWER) control, examining the correlation between a phenotype of interest with the genotype of genetic variants one at a time, while also accounting for non-genetic risk factors and covariates. We refer to this as the conventional GWAS pipeline in this paper. Generalized linear models allow for the analysis of non-Gaussian phenotypes, such as dichotomized disease status, and mixed effect models account for population structure and polygenic effects.

Although this approach has yielded a large number of SNP-phenotype associations,^6^ two problems remain open and are becoming a crucial focus of the post-GWAS era. First, growing evidence supports the highly polygenic nature of many complex traits. To capture a substantial amount of heritability and construct reliable predictive models, it is necessary to use a multitude of loci, each with small effects, going well beyond the set of variants that are identified as significant in GWAS based on stringent adjustment for multiple comparisons that control the FWER.^7,8^ This is reflected in the common rules to construct polygenic risk scores with more liberal p-value cutoff and explains the success of prediction rules based on methods like the lasso, which offers no control on the selection of variables.^9^ Second, the association findings from standard GWAS are too repetitive and imprecise, hampering the transition from genetic discoveries to biological mechanisms. Traditional marginal tests identify any proxy feature correlated with the true causal variants. Second-stage fine-mapping approaches then focus the researcher’s attention on smaller sets of variants which have some guarantee of harboring the causal variant with high probability.^10,11^ However, these methods are inherently constrained by their focus on strong associations detected with marginal models; they cannot lead to discoveries of causal variants at other loci, nor provide genome-wide error control guarantees. Additionally, when multiple credible sets are identified at the same locus, these do not unequivocally represent independent causal effects. This ambiguity is particularly relevant given recent findings suggesting that genetic association signals at a locus can be the result of multiple causal variants.^12^

In contrast to marginal models with FWER control, recent work has shown that searching for conditional independent effects with False Discovery Rate (FDR) control can enhance the detection of variants with weaker effect sizes and improve the chances of identifying putatively causal variants.^13^ A conditional independence hypothesis evaluates the effect of each genetic variant on a phenotype, conditioning on all other genetic variants throughout the genome. The knockoffs methodology is a recently proposed statistical framework for testing these conditional independence hypotheses in high-dimensional settings.^14^ This approach involves generating synthetic, noisy replicas (knockoffs) of the original genetic variants, which function as negative controls for the conditional tests. The knockoffs aid in the selection of significant genetic variants and help mitigate the confounding effect of linkage disequilibrium (LD).^15–17^ Several knockoffs-based methodologies have been proposed for genetic research.^13,14,16,18,19^ The variants identified by the knockoff approach were shown more likely to be causal. The connections between conditional testing and causal inference have been further analyzed for genetic trio studies and in studies searching for consistent conditional associations across environments.^20,21^ Motivated by the frequent unavailability of individual-level data in large meta-analyses of GWAS, He et al. (2022) developed GhostKnockoffs, which requires only summary statistics and enables a knockoff approach without access to individual genotypes.^22^ However, although testing for conditional independent hypothesis, GhostKnockoffs defines marginal Z-scores as feature importance, where the statistical power can be suboptimal compared to using feature importance from modern machine learning methods that better characterize the conditional independent effect. Moreover, there are still considerable challenges in performing computationally efficient conditional independence tests in the context of large-scale GWAS.

In this study, we introduce an analytical pipeline for genome-wide detection of putative causal variants, which integrates several technical advancements described in detail in a series of companion papers^23,24^ and allows, for the first time, to seamlessly wrap a knockoff analysis and a genome-wide sparse regression around any GWAS results. Without changing any of the GWAS processing steps, our pipeline enhances the discovery of additional loci and localizes conditionally independent causal effects. We define the causal effect of a genetic variant, in a manner that is conceptually equivalent to quantifying the change in phenotypic value observed by introducing a sequence change in a functional experiment, such as massively parallel reporter assay (MPRA) or CRISPR-Cas9. To achieve this, we rely on several technical advancements, as (1) the ability to use summary statistics (e.g., p-values and direction of effects), from potentially overlapping studies, to jointly assess all genetic variants across the genome via ultrahigh-dimensional sparse regression with FDR control. The new approach with ultrahigh-dimensional sparse regression is substantially more powerful than the original version of GhostKnockoffs developed by He et al. (2022), which use marginal test statistics as feature importance;^24^ (2) the optimized construction of group knockoffs to enhance the power to identify tightly linked causal variants.^23^ We implement the approach in a user-friendly open-source software, which analyzes genome-wide summary statistics on a single central processing unit (CPU) in less than 15 minutes.

We illustrate performance via simulations and by applying our new analysis pipeline to a meta-analysis of ten large-scale genetic studies of Alzheimer’s disease (AD) since 2017. This analysis identified 82 loci associated with AD, including 37 additional loci missed by the conventional GWAS pipeline via marginal association tests. We used data on MPRA + CRISPR-Cas9 experiments to validate these identified variants and to show that the identified putative causal variants achieve good agreement with the experimental results. Furthermore, using functional genomics data from single-cell assays for transposase-accessible chromatin using sequencing (scATAC-seq) in excitatory neurons, inhibitory neurons, microglia, oligodendrocytes, astrocytes and OPCs, we show functional enrichment of these variants in microglia. Finally, to demonstrate the generalizability of the method, we apply the method to summary statistics from a collection of large-scale GWAS for a variety of phenotypes. On average, the method identifies 22.7% more loci and 78.7% fewer proxy variants per locus compared to the conventional GWAS pipeline via marginal association tests. The results highlight the appealing attributes of the proposed method for robustly uncovering additional loci and pinpointing putative causal variants that underlie each locus. We have made the discoveries and software freely available to the community and anticipate that routine end-to-end in silico identification of putative causal genetic variants will become an important tool that will facilitate downstream functional experiments and future research into disease mechanisms and potential therapies.

## Results

### Overview of the method: causal inference of genetic variants to attenuate LD confounding

We assume a study population of *n* independent individuals, on which we measure *p* genetic variants ***G*** = (*G*_1_, …, *G*_*p*_) and a phenotype *Y*. In conventional genome-wide association studies, we test the null hypothesis *H*_0_: *Y* ⊥ *G*_*j*_ for each genetic variant *j*, referred to as marginal association testing. It is essential to note that, without randomization-based assignment of *G*_*j*_ to level *g*_*j*_, the usual marginal association between *Y* and *G*_*j*_ commonly assessed in GWAS cannot uncover causation. One major confounding effect in genetic studies is linkage equilibrium (LD), whereby a non-causal variant can be identified by a marginal association test if it is correlated with a causal variant. To evaluate the causal effect of the *j*-th genetic variant on *Y*, we are interested in testing the following conditional independence (CIT) hypothesis:

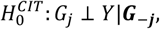

where ***G***_−***j***_ = (*G*_1_, …, *G*_*j*−1_, *G*_*j*+1_, …, *G*_*p*_) are all variants across the genome except the *j*-th. The CIT hypothesis has a causal inference interpretation. Under standard identifiability conditions in causal inference, we show in **Materials and Methods** that this hypothesis is equivalent to testing whether a nonparametric conditional causal effect (CCE) in causal inference is zero,

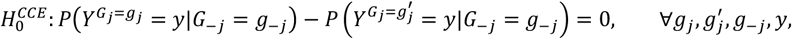

where the causal effect of *G*_*j*_ on *Y* is defined on the basis of the counterfactual (or potential) outcomes 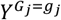 and 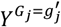, which are mutually unobservable quantities.^25,26^ Intuitively, the CCE evaluates the effect of *G*_*j*_ on *Y* by considering changing the value of *G*_*j*_ from *g*_*j*_ to 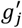, without altering any other variants. If the distribution of *Y* changes for any stratum *G*_−*j*_ = *g*_−*j*_, the *j*-th variant may be considered to have a causal effect on *Y*. We adopt this measure of causal effect because it is conceptually similar to a functional experiment (e.g., MPRA or CRISPR) that edits a particular sequence and then looks for any change in the phenotype. One of the main conditions is the unconfoundedness, which assumes no unmeasured confounders beyond *G*_−*j*_. In the context of attenuate confounding effects induced by linkage disequilibrium, this means that the causal effect can be inferred if all genetic variants are genotyped. A more detailed description of the method and discussion about the connections with other measures of causal effect is included in **Materials and Methods**.

Unlike the usual two-stage marginal association test + fine-mapping procedure, the conditional independence testing in a high-dimensional regression model simultaneously performs discovery and prioritization of causal variants with improved power and reduced LD confounding as demonstrated in Sesia et al. (2020).^13^ In this paper, we show how summary statistics (e.g., p-values and direction of effects) can be leveraged to achieve these appealing properties. The ability to use summary statistics facilitates the integration of multiple studies to the maximum extent feasible, and it allows the method to be flexibly applied on top of the standard GWAS pipeline without changing any of the processing steps.

### Numerical Experiments and Simulations

We performed simulations to empirically evaluate the performance of the proposed conditional independence tests in terms of discovering and mapping causal variants.

For variant discovery, we evaluate type I error rate (false discovery rate; FDR) and power based on the average of 500 replicates, defines as follows:

- Type I error: the percentage of identified variant that are not in the same LD group of any causal variant.
- Power: the percentage of causal variants that are discovered.

The primary comparison methods include: 1) The proposed conditional independence test. We implemented two versions, depending on how the feature importance is defined – one based on a Lasso-type genome-wide sparse regression (CIT-Lasso), and the other one based on separately applying Sum of Single Effects (SuSiE) model to every LD block across the genome (CIT-SuSiE); 2) The original GhostKnockoffs developed by He et al. (2022) on the basis of marginal Z-scores (CIT-Marginal); 3) Marginal association tests with Benjamini-Hochberg adjustment for FDR control (MAT-BH); 4) Marginal association tests with matched number of discoveries made by CIT-Lasso (MAT-Matched). That is, we rank variants by marginal test p-values and select top variants, where the number is set to be same as the discoveries made by CIT-Lasso. We are looking for a method with higher power while the FDR is under the target level (0.1).

Turning to fine mapping, we evaluate three metrics of interest:

- Size: number of variants in the catching/credible sets; we report the maximum and average size of all identified catching/credible sets for each replicate.
- Purity: the smallest squared correlation among all pairs of variants within a catching/credible set; we report the minimum purity and average purity of all identified catching/credible sets for each replicate.
- Fine-mapping power: the probability that the causal variant is covered by the catching/credible sets for the discovered loci.

While the average size and purity reflect the average performance, the maximum size and minimum purity reflect the performance for more challenging tasks. The primary comparison methods include: 1) and 2) as above; 3) SuSiE fine-mapping based on reported credible sets. The SuSiE credible sets are based on the default setting in the susieR package, with the maximum number of non-zero effects in the SuSiE regression model equal to 10, coverage equal to 95% and minimum absolute correlation allowed in a credible set equal to 0.5. It is worth noting that SuSiE was not developed for discovery. To assure that approaches are compared on a common ground, we apply SuSiE to the same loci discovered by CIT-Lasso. We are looking for a method with smaller catching/credible sets, higher purity and higher fine-mapping power.

The genetic data in our simulations is constructed from the UKbiobank. The simulation study contains 500 replicates. For each replicate, we sampled 15,000 individuals with real genotype data on variants with minor allele frequency (MAF) ≥0.01. We sample 200kb regions from 500 nearly independent LD blocks (one region per block) where the 500 blocks account for ∼30% of the genome and the selected 200kb regions account for ∼3.5% of the genome. We restricted the simulations to variants with MAF ≥0.01 to ensure stable calculation of summary statistics (e.g. p-values). We used 5,000 of the 15,000 individuals as the reference panel to estimate the LD structure and considered the remaining 10,000 individuals as part of the target study to compute the Z-scores. We considered a relatively small sample size for the reference panel to demonstrate the potential application of the proposed method to underrepresented populations, where the current sample sizes are substantially smaller (e.g. the Pan-UKBB panel has 420,531 samples for European ancestry but only 6,636 samples for African ancestry).

To simulate the trait, we randomly set 50 regions to be causal, each with one directly genotyped causal variant. The quantitative trait and dichotomous trait are then simulated as follows, respectively:

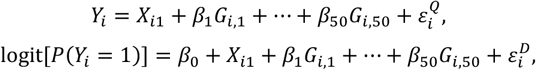

where 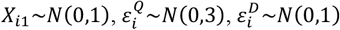 and they are independent; *X*_01_ is the observed covariate that is adjusted in the analysis; 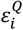 and 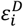 reflect variation due to unobserved covariates; (*G*_0,1_, …, *G*_0,20_) are selected risk variants. We set the effect 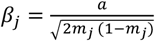, where *m*_*j*_ is the MAF for the *j*-th variant. We define *a* such that the variance due to the risk variants, 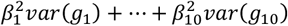, is 1.2 for quantitative trait and 1.5 for dichotomous trait; For dichotomous trait, we set *β*_0_ such that the disease prevalence is 0.1. We performed marginal score test for each single variant to compute Z-scores, adjusting for the observed covariate *X*_*i*1_.

### Variant discovery: type I error and power

**Figure 1** shows that the proposed method CIT-Lasso and its alternative SuSiE-type implementation, CIT-SuSiE, exhibits substantially higher power than the original GhostKnockoffs (He et al. 2022; CIT-Marginal) where the test statistics are on the basis marginal Z-scores. The higher power is because the joint modeling of the effects of all variants avoids comparing the signal of each individual variant with a “large” error term, which encapsulates the combined effect of all other variants. Also, the test statistics defined by a sparse regression better characterizes the conditionally independent effect on the outcome variable compared to the marginal test statistics. All methods have valid FDR control at the target level, except MAT - BH. This is substantially due to the fact that there is a discrepancy between how marginal tests count false/true discoveries and the assessment one makes of findings from a casual viewpoint, which is the one we take here. From the marginal p-values point of view proxy variants that are only correlated with the causal ones are true positives. This results in a large number of possible true rejections, which, in turn, in an adaptive method like the BH, results in the lowering of the threshold for significance.^27^ The false discoveries that this opens the door to, typically, are spread across multiple loci: through the causal lenses, this disproportionally increases the number of false rejections. MAT with matched number of discoveries (MAT-Matched) represents an attempt to remedy this lack of FDR control. However, it leads to lower power compared to CIT methods. This is because MAT makes multiple repetitive discoveries from the same LD group: with the same number of total discoveries MAT-Matched can cover only a fraction of the loci identified by CIT-Lasso and CIT-SuSie. The results highlight the fact that FDR control in GWAS is particularly challenging, as previously discussed by Sesia et al. (2020).

### Fine-mapping metrics

**Figure 2** shows that the proposed methods CIT-Lasso and CIT-SuSiE catching sets have substantially smaller size and higher purity compared to CIT-Marginal and SuSiE credible sets. The improvement over CIT-Marginal is expected and likely due to the fact that the proposed method utilizes a sparse regression to enhance the prioritization of causal variants, as opposed to measuring importance with marginal Z-scores. More striking is the improvement in size and purity over SuSiE. In addition, the proposed methods, including the CIT-SuSiE, lead to catching sets with substantially smaller size and higher purity than the credible sets defined by the original implementation of SuSiE. The results demonstrate a difference in the way catching/credible sets are constructed. While SuSiE’s defines credible sets based on coverage, the catching set defined by CIT-Lasso or CIT-SuSiE controls false discovery rate of the identified variants. More importantly, we observed that CIT-Lasso and CIT-SuSiE exhibit similar or better fine-mapping power compared to SuSiE credible set while the size and purity of catching/credible sets are much improved.

**Figure 1:**
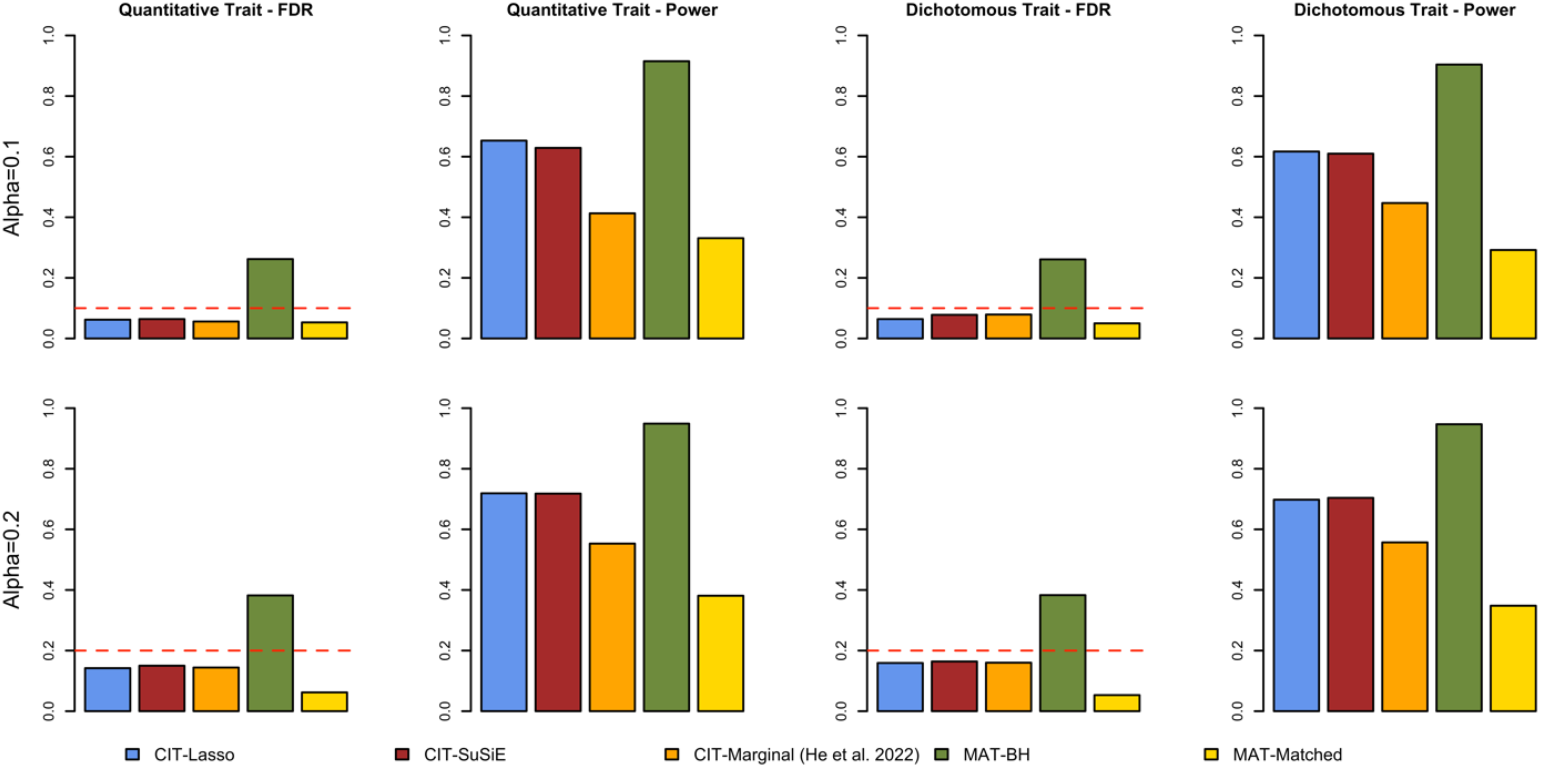
Genome-wide simulation study – FDR and power. The simulation study is based on 500 replicates. Each replicate includes 500 approximately independent 200kb regions across the genome, among which 50 contain one causal variant. CIT-Lasso/SuSiE: the proposed conditional independence test with importance measure derived from a Lasso type model / SuSiE type model. CIT-marginal: the original GhostKnockoffs by He et al. (2022), where the feature statistics are defined by marginal tests. MAT-BH: marginal association test with Benjamini-Hochberg correction for FDR control. MAT-Matched: we rank variants by marginal test p-values and select top variants, where the number is set to be same as the discoveries made by CIT-Lasso.

**Figure 2:**
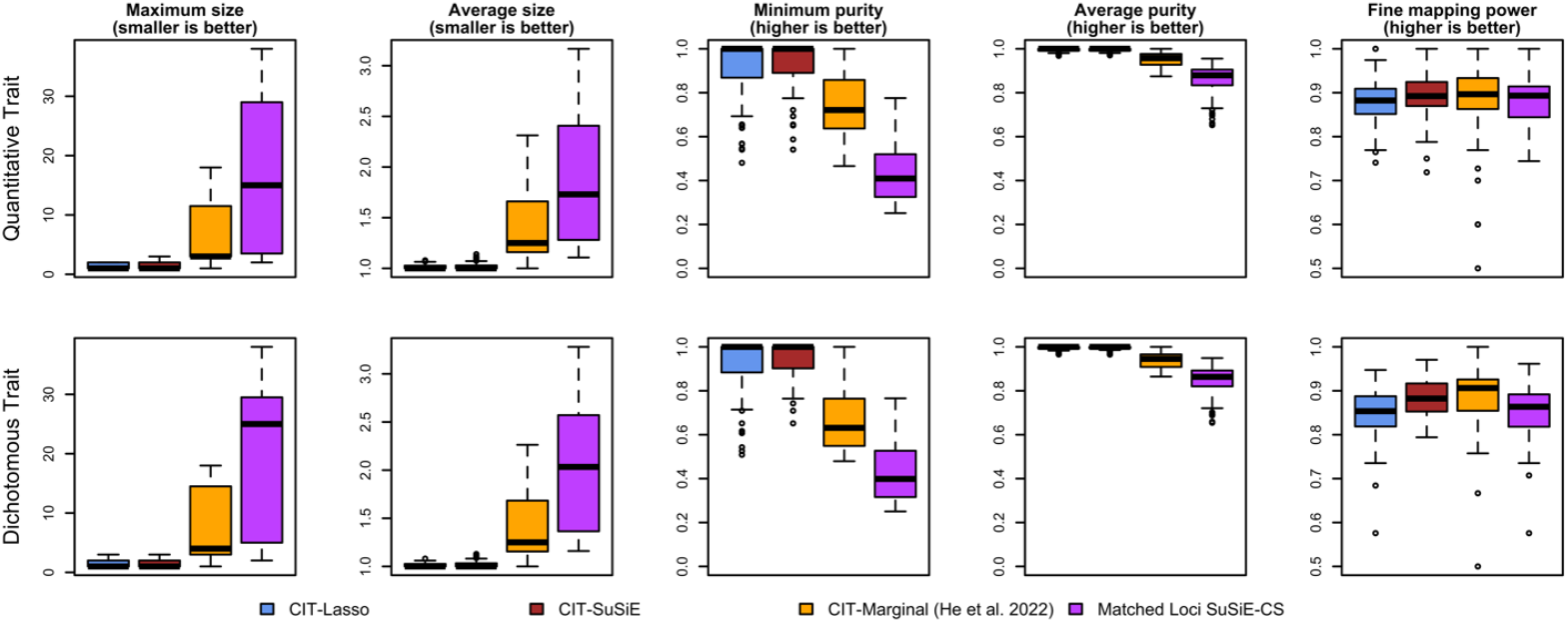
Genome-wide simulation study – mapping causal variants. CIT-Lasso/SuSiE: the proposed conditional independence test with feature importance from Lasso type model / SuSiE type model. CIT-marginal: the original GhostKnockoffs by He et al. (2022), where the feature statistics are defined by marginal tests. Matched Loci SuSiE-CS: the credible sets reported by SuSiE. It is worth noting that SuSiE was not developed for discovery. To make fair comparison, we apply SuSiE to the same loci discovered by CIT-Lasso.

### Application to Alzheimer’s disease genetics

Alzheimer’s disease (AD) is the most common cause of dementia among people over the age of 65, affecting an estimated 5.5 million Americans. To study genetic risk and to identify molecular mechanisms of AD, we applied the proposed method to summary statistics from ten overlapping large-scale array-based genome-wide association studies and whole-exome/-genome sequencing studies since 2017, including: 1. The genome-wide survival association study by Huang et al. (2017) (14,406 cases, 25,849 controls); 2. The genome-wide meta-analysis by Jansen et al. (2019) (71,880 clinically diagnosed/proxy AD cases, 383,378 controls); 3. The genome-wide meta-analysis by Kunkle et al. (2019) (21,982 cases, 41,944 controls); 4. The genome-wide meta-analysis by Schwartzentruber et al. (2021), aggregating Kunkle et al. (2019) and UK Biobank based on a proxy AD phenotype; 5. An in-house genome-wide association study of 15,209 cases and 14,452 controls aggregating 27 cohorts across 39 SNP array data sets, imputed using the TOPMed reference panels; 6-7. Two whole-exome sequencing analyses of data from The Alzheimer’s Disease Sequencing Project (ADSP) by Bis et al. (2019) (5740 cases, 5096 controls), and Le Guen et al. (2021) (6008 cases, 5119 controls); 8. An in-house whole-exome sequencing analysis of ADSP (6155 cases, 5418 controls); 9. An in-house whole-genome sequencing analysis of the 2021 ADSP release (3584 cases, 2949 controls); 10. The genome-wide meta-analysis by Bellenguez et al. (2022) (discovery phase; 85,934 clinically diagnosed/proxy AD cases and 401,577 controls).^28–36^ All studies focused on individuals with European ancestry. We used LD matrices estimated via the Pan-UK Biobank data.^37^ We restricted the analyses to directly genotyped common and low-frequency variants with minor allele frequency >1%.

We propose a meta-analysis strategy that adaptively estimates weights to combine the ten studies, allowing for sample overlap, as described in the **Appendix**. The meta-analysis Z-scores serve as the input of the proposed method. We present the estimated study correlations and the estimated optimal weights in **Figure 3**. The correlation results are consistent with our knowledge of overlap and other factors, such as differences in phenotype definition, analysis strategies, and quality control. The corresponding scheme up-weights studies that are large in size and carry independent information and down-weights studies that largely overlap with others. Notably, the three major AD genetic studies, Jansen et al. (2019), Schwartzentruber et al. (2021) and Bellenguez et al. (2022), are estimated to have larger weights compared to the other studies which were integrated into the three meta-analyses.^29,31,32^ In addition to the meta-analysis, we applied the method separately to summary statistics from the three major AD genetic studies. We present the main meta-analysis results as a Manhattan plot in **Figure 4**. (the corresponding display for conventional GWAS meta-analysis is in **Supplementary Figure 1)**. We define two loci as different if they are at least 1Mb away from each other, where each locus may contain one or multiple sets of putative causal variants with conditionally independent effects. We adopt the most proximal gene’s name as the locus name, recognizing that it is not necessarily the causal gene.

**Figure 3:**
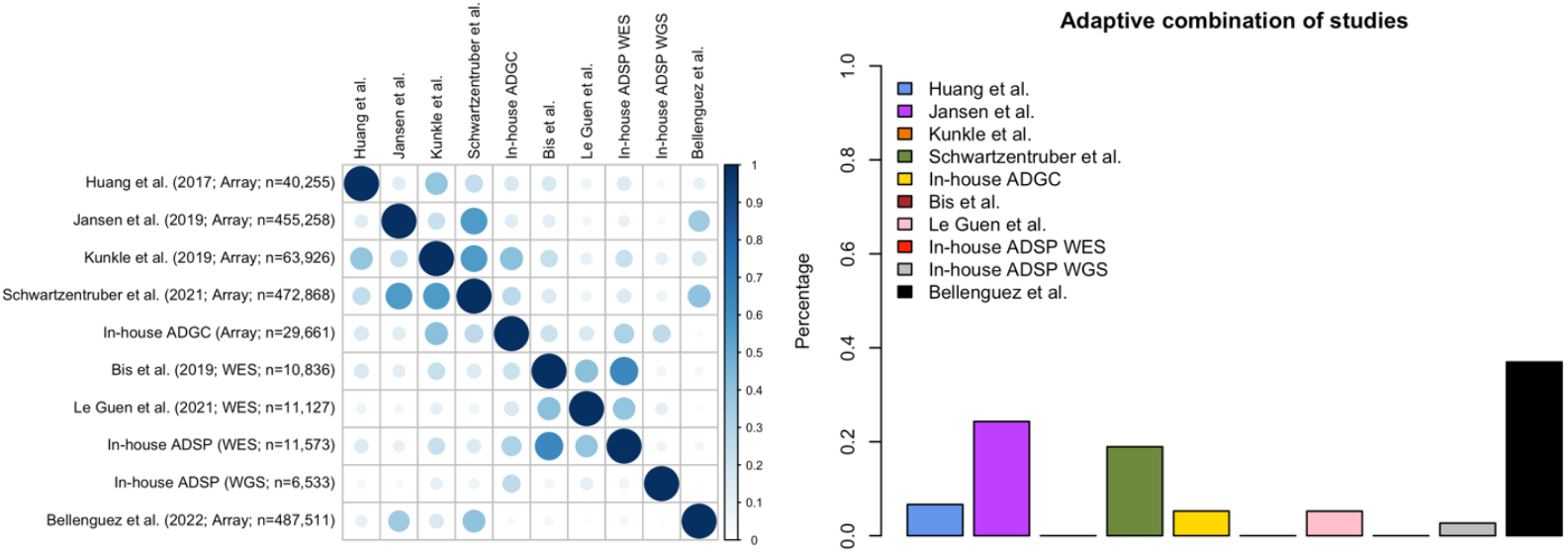
Study design. Our analysis of Alzheimer’s disease genetics is an aggregation of ten possibly overlapping studies since 2013. Left panel: estimated study correlations; For each study, we present genotyping technology and sample size. Right panel: estimated adaptive combination of studies; studies with larger sample size and less correlation with other studies have larger weights. Each bar presents the weight per study in percentage, i.e., weight per study divided by the summation of all weights.

**Figure 4:**
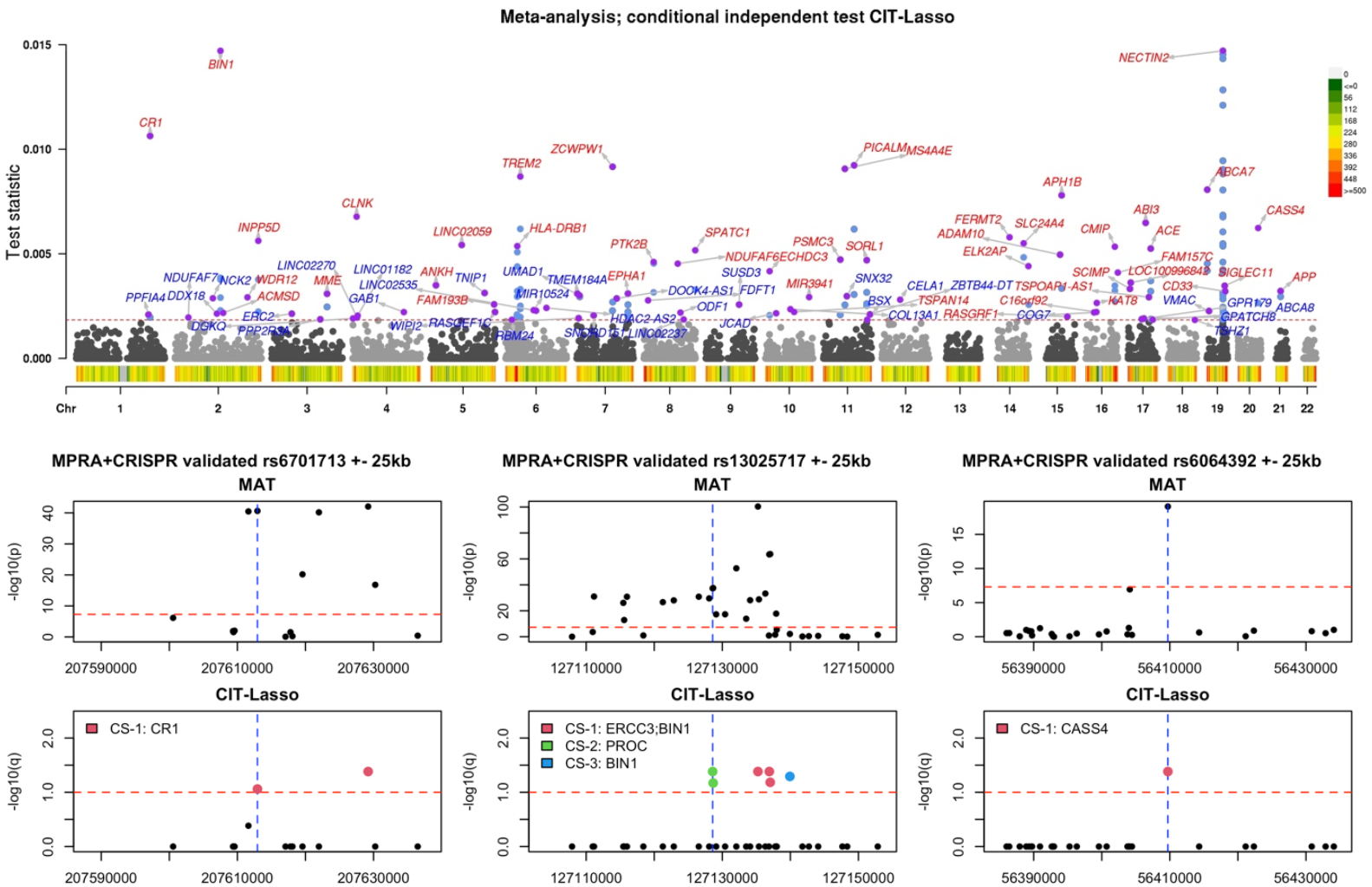
Meta-analysis of Alzheimer’s disease genetic studies. Top panel: Manhattan plot from the proposed conditional independence tests (CIT-Lasso). The dotted lines indicate the significant threshold corresponding to FDR=0.10. Two loci are considered distinct if they are at least 1Mb away from each other. For each locus, we annotated the variant with the largest W statistic and adopt the most proximal gene’s name as the locus name. Loci that are less than 1Mb away from loci (*p* ≤ 5 × 10^56^) idenitified with a conventional GWAS pipeline are highlighted in red (45 loci). Additional loci identified by CIT-Lasso are highlighted in blue (37 loci). Variant density is shown at the bottom of plot (number of variants per 1Mb). Bottom panels: we present fine-mapping examples for rs6701713 (*CR1*; MPRA *p* = 8 × 10^522^; CRISPR-Microglia *p* = 2.9 × 10^52^), rs13025717 (*BIN1*; MPRA *p* = 2.4 × 10^589^; CRISPR-Microglia *p* = 1.3 × 10^5:^) and rs6064392 (*CASS4*; MPRA *p* = 1.04 × 10^5;;^; CRISPR-Microglia *p* = 4.2 × 10^5;^), three causal variants validated by Cooper et al. (2022) using massively parallel reporter assays (MPRA) coupled with CRISPR in neurons and microglia. We compare marginal association test (MAT) and CIT-lasso. Different colors represent different catching sets (for CIT-Lasso, they represent independent conditional causal effects). The legend presents the genes that each catching set potentially regulates, mapped by the cS2G method. The red dotted lines represent the p-value threshold of 5 × 10^56^ and FDR threshold of 0.10 for MAT and CIT-lasso respectively. The blue dotted lines represent the location of the validated causal variants.

We observed that CIT-Lasso (target FDR=0.1) consistently identified more loci compared to marginal association tests as performed in conventional GWAS (35 loci vs. 23 loci for Jansen et al. (2019); 31 loci vs. 28 loci for Schwartzentruber et al. (2021); 68 loci vs. 42 loci for Bellenguez et al. (2022); 82 loci vs. 51 loci for our meta-analysis; results are summarized in **Supplementary Figure 2**). Meanwhile, the conditional independence test results in a substantially smaller number of proxy variants per locus (on average, 3.3 vs. 19.3 for Jansen et al. (2019); 1.6 vs. 18.8 for Schwartzentruber et al. (2021); 2.3 vs. 15.8 loci for Bellenguez et al. (2022); 2.6 vs. 18.8 for our meta-analysis). Overall, the results demonstrate that the proposed method is able to robustly discover additional loci that are missed by marginal association test, and at the same time pinpoint potential causal variants underpinning each locus as a fine-mapping method would.

### Concordance with MPRA+CRISPR experiments

To additionally evaluate the validity of the identified putative causal variants, we leveraged data and results from Cooper et al. (2022), which used MPRA to screen noncoding variants reported in previous AD GWAS, followed by CRISPR functional validation in neurons and microglia.^38^ In this experimental approach, the functional effect of a variant is measured by evaluating the effect on gene expression of the corresponding sequence alteration. This is seen as the gold-standard approach for validating functional consequences of a variant (referred to as a causal variant in this section). We evaluated how the variants identified by CIT-Lasso overlap with the nine MPRA+CRISPR validated variants (eight variants reported by Cooper et al. (2022) and one additional variant with p-value ≤ 0.05 in both MPRA and CRISPR experiments), and additionally compared results with SuSiE.

Among the nine causal variants, three are directly genotyped and all of them are correctly identified by CIT-Lasso as putative causal variants. We present them in **Figure 4**, and they correspond to gene *CR1, BIN1* and *CASS4*. For the *CR1* locus, rs6701713 is a causal variant validated by MPRA (*p* = 8 × 10^−33^) and CRISPR (microglia; *p* = 2.9 × 10^−3^). For the *BIN1* locus, rs13025717 is a causal variant validated by MPRA (*p* = 2.4 × 10^−41^) and CRISPR (microglia; *p* = 1.3 × 10^−2^). For the *CASS4* locus, rs6064392 is a causal variant validated by MPRA (*p* = 1.04 × 10^−22^) and CRISPR (microglia; *p* = 4.2 × 10^−2^).

Although SuSiE is not designed for genome-wide discovery of causal variants as CIT-Lasso does, we compared their fine-mapping performance in pinpointing the causal variant for the same region and present the results in **Supplementary Figure 3**. For the *CR1* locus, we observed that SuSiE is able to include the functional variant as part of its credible set, but the PIP is close to 0 and we cannot determine whether it should be selected. CIT-Lasso is able to provide a selection set that empirically controls the FDR. For the *BIN1* locus, the functional variant is not covered by any credible sets identified by SuSiE, while CIT-Lasso correctly identified it as a putative causal variant. For the *CASS4* locus, both SuSiE and CIT-Lasso are able to pinpoint the causal variant.

The other six causal variants are not directly genotyped and therefore are not present in our primary analysis. We present them in **Supplementary Figure 4**. We found that CIT-Lasso can identify tightly linked neighboring variants as the causal variants in some cases. For instance, rs1532277 is a causal variant validated by MPRA (*p* = 1.5 × 10^−>^) and CRISPR (Neuron; *p* = 6.4 × 10^−2^) at the *CLU* locus. While rs1532277 is not directly genotyped in the dataset we considered, CIT-Lasso identify its tightly linked neighbor rs1532278 (*D*^/^ = 1; 134 base pairs away).

### Variants with weaker effect sizes solely identified by CIT-Lasso are replicated and functionally enriched

Since the proposed meta-analysis aims to incorporate all major studies to date, which are possibly overlapping with each other, we do not have a hold-out independent dataset for replication. To validate the proposed method, we adopted three alternative strategies: 1. We investigated whether the variants identified in an earlier study (e.g., Jansen et al. 2019) were replicated with smaller p-values in a future study with increased sample size (e.g., the meta-analysis); 2. We evaluated whether the identified variants were more likely to be functional compared to genome background variants; 3. We confirmed that variants identified by the meta-analysis exhibited consistent associations across the ten studies. We are particularly interested in performing these validation strategies on the additional loci solely identified by the proposed method but missed by marginal association tests.

We first evaluate whether the variants identified by our proposed method can be replicated in a future larger study with smaller marginal p-values. We compared the distribution of p-values in the meta-analysis with those of Jansen et al. (2019) and Schwartzentruber et al. (2021), respectively. We present the results in **Figure 5**. We observe that the p-values of the variants identified in previous studies become generally smaller in the meta-analysis with an increased sample size (**Figure 5**, left panel). For variants from the additional loci solely identified by our proposed method but missed by conventional GWAS (*p* ≥ 5 × 10^−8^), a large proportion of them are identified as statistically significant in the meta-analysis (**Figure 5**, right panel; 39% for Jansen et al. 2019; 60% for Schwartzentruber et al. 2023). This demonstrates the power of the proposed method, which allows identification of the weaker associations in earlier studies with smaller sample sizes.

**Figure 5.**
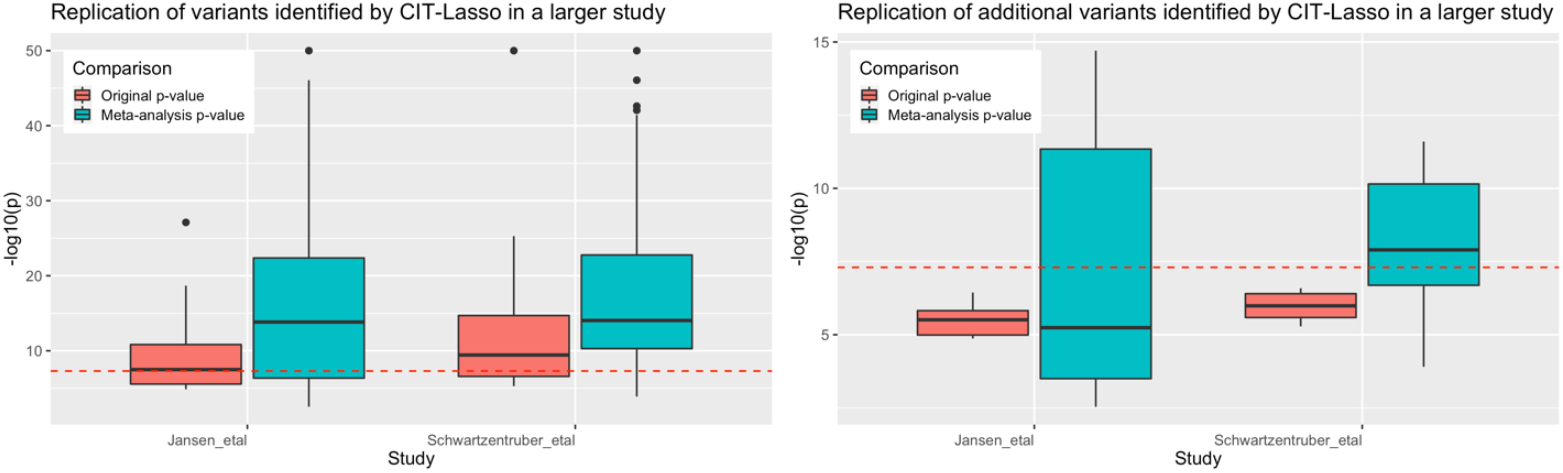
Variants identified by CIT-Lasso are replicated in a larger study with smaller marginal p-values. For variants identified by CIT-Lasso, we compare the marginal p-values in the original study and the marginal p-values in the meta-analysis. Results show that a large proportion of variants identified by CIT-Lasso but missed by the conventional marginal association test (loci that are more than 1Mb away from conventional GWAS loci) reach genome-wide significance level in a future study with larger sample size (38.9% for Jansen et al. 2019; 60.0% for Schwartzentruber et al. 2021). We excluded *APOE* locus and truncated p-values at 10^5:3^ for better visualization.

Second, we investigate whether the identified variants are functionally enriched. We leveraged results from single-cell assay for transposase-accessible chromatin using sequencing (scATAC-seq) in excitatory neurons, inhibitory neurons, microglia, oligodendrocytes, astrocytes and OPCs.^39^ We evaluated how our identified variants overlap with the scATAC-seq peaks identified by Corces et al. (2020). In **Figure 6**, we present the enrichment relative to the background genome, defined as

**Figure 6.**
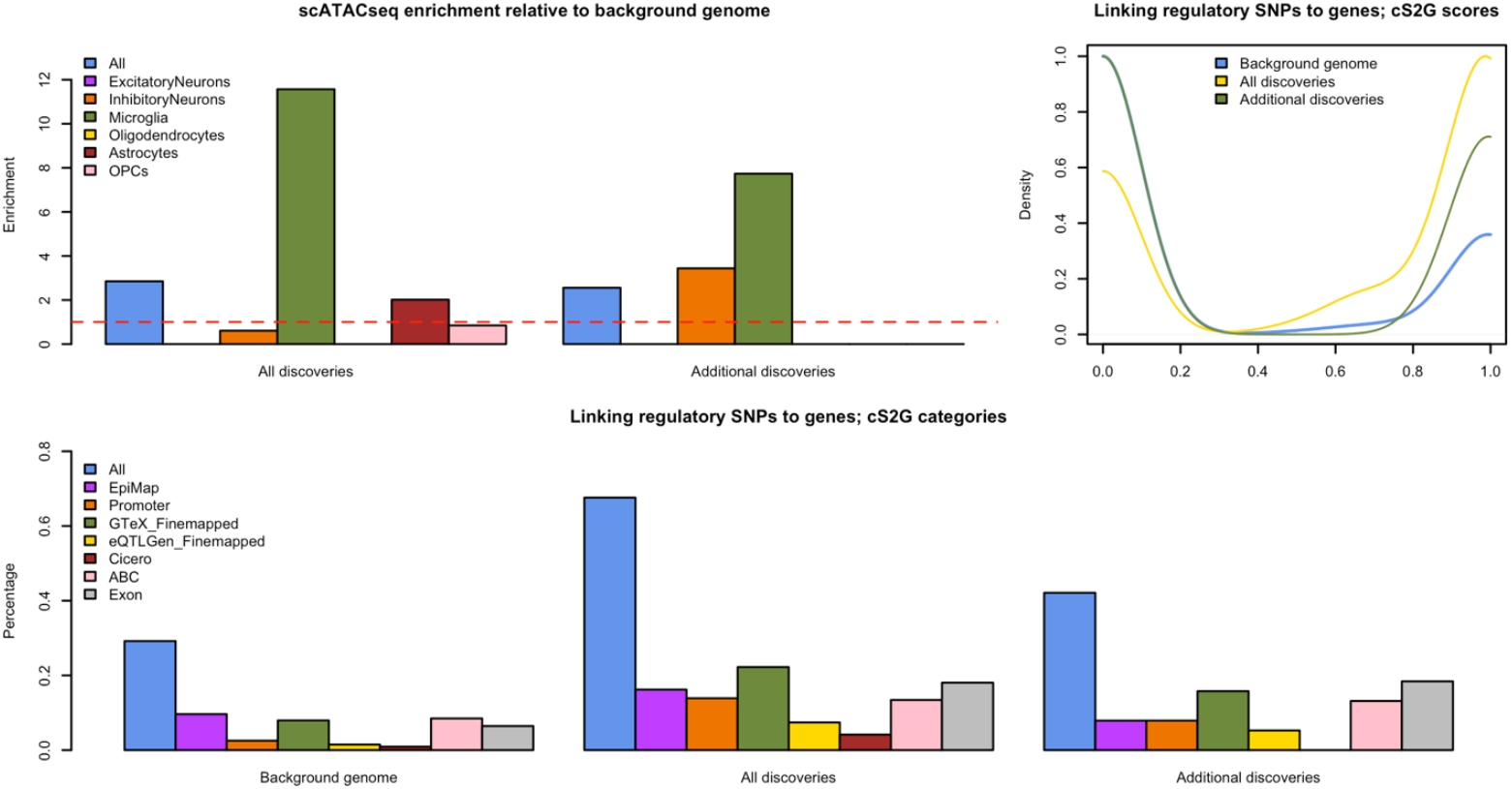
Variants identified by CIT-Lasso are functionally enriched. The additional variants identified by CIT-Lasso have similar level of enrichment. Top-left panel: variants identified by CIT-Lasso are more likely to overlap with scATACseq peaks in microglia. The enrichment is calculated as proportion of identified variants that overlap with scATACseq peaks divided by that of the background genome. Top-right panel: variants identified by CIT-Lasso have higher cS2G score to be functionally mapped to a gene that they potentially regulate. The figure presents the distribution of cS2G scores of the identified variants relative to that of all background genome variants. Bottom panel: variants identified by CIT-Lasso are more likely to be in a cS2G functional category.

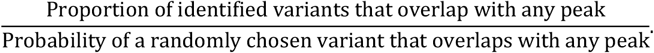

We observed that the identified variants exhibit strong enrichment (>10x) in microglia relative to the background genome. Interestingly, the additional variants identified by the proposed method, but missed by conventional GWAS (≥ 5 × 10^−8^), have similar levels of enrichment in microglia compared to those exhibiting stronger associations.

We also leveraged a recent combined SNP-to-gene linking strategy (cS2G) proposed by Gazal et al. (2022) to annotate the identified variants and map them to the genes they potentially regulate.^40^ The cS2G method combined seven S2G functional categories, including exon, promoter, GTEx fine-mapped cis-eQTL, eQTLGen fine-mapped blood cis-eQTL, EpiMap enhancer-gene linking, Activity-By-Contact (ABC), scATAC-seq Cicero blood/basal. A cS2G linking score is then computed to summarize the functional consequences and the confidence of the gene mapping. In **Figure 6**, we present the distribution of the cS2G linking score and the proportion of variants that fall in each of the functional categories. We observed that variants identified by the proposed method have substantially higher cS2G linking scores relative to genome background variants. They also have substantially higher probability of belonging to one of the functional categories. Similar enrichment is found for the additional variants solely identified by the proposed method. The results show that the proposed method can identify putatively functional variants with weaker effects that are missed by conventional association tests.

Finally, we checked whether the variants solely identified by the proposed method but missed by conventional marginal association tests exhibit concordant Z-scores and directions of effect across the ten studies. We calculated the Spearman correlation of Z-scores between each pair of studies involved in the meta-analysis. For each identified variant, we also computed the proportion of studies where the direction of effect is concordant with that of the meta-analysis. We present the results in **Supplementary Figure 5**. We observed that Z-scores of the identified variants are positively correlated across the ten studies (median correlation = 0.7). In addition, almost all identified variants have concordant directions of effect when compared to the meta-analysis (for 94% variants, the direction of effect in the meta-analysis is concordant with >80% individual studies). This demonstrates that the proposed method, paired with the meta-analysis strategy, can robustly identify genetic variants that share consistent association across the ten AD genetic studies since 2017.

### Generalizability of the method: application to large-scale GWAS summary statistics

The proposed method simultaneously produces discovery and prioritization of causal variants in 15 minutes with only a single CPU. This opens the exciting possibility of identifying putative causal variants at the phenome-scale and beyond. In this section, we demonstrate the generalizability of the proposed method by applying it to summary statistics from large-scale GWAS since 2013. Specifically, we curated GWAS summary statistics from 400+ publications. We restricted our analysis to studies with sample size >100,000, with individuals of European ancestry and at least ten loci passing the genome-wide significance level (marginal p-value ≤ 5 × 10^−8^), to focus on traits with appreciable polygenicity. This results in 67 studies. We applied CIT-Lasso to the corresponding summary statistics and compared the results with the original GWAS via MAT; we compared the number of discovered loci and the average number of proxy variants per locus. We additionally evaluated the proportion of studies where the proposed method identifies more loci than marginal association tests as a function of phenotype polygenicity (quantified by the number of loci discovered by the original GWAS). We present the results at FDR=0.1 and 0.2 in **Figure 7**.

**Figure 7.**
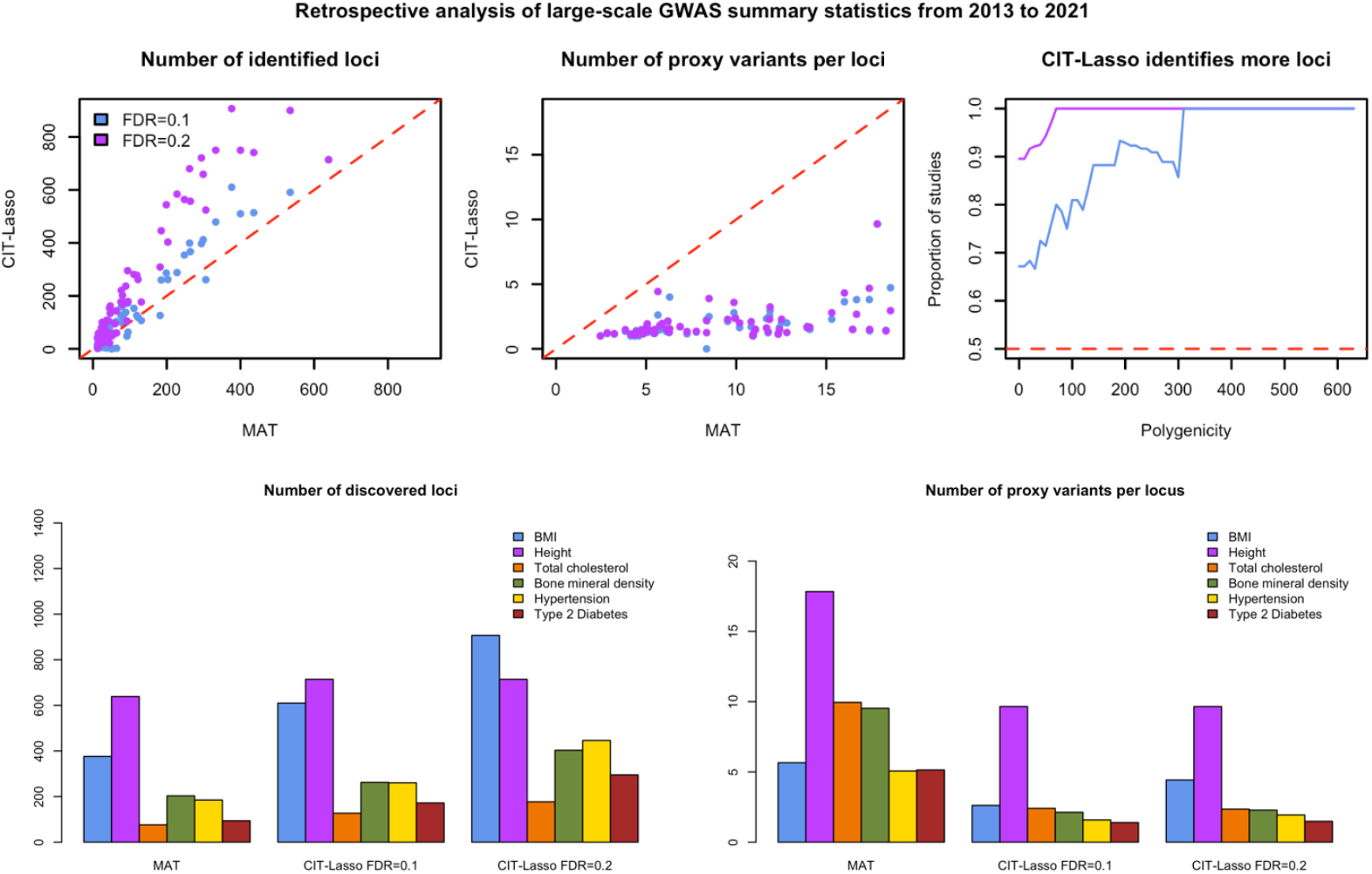
Retrospective analysis of large-scale GWAS summary statistics since 2013. Left panel: number of loci identified by CIT-Lasso vs. marginal association test (MAT); Mid panel: average number of proxy variants per loci by CIT-Lasso vs. MAT; Right panel: proportion of studies that CIT-lasso identifies more loci than MAT among all studies with a polygenicity of the trait greater than a particular level. The polygenicity is quantified by the number of loci identified by MAT as in the original GWAS.

We observe that CIT-Lasso generally identifies a substantially smaller number of proxy variants per locus compared to MAT (top mid panel). On average (over the 67 studies), CIT-Lasso identified 1.83 proxy variants whereas MAT identifies 8.61 proxy variants (down 78.7%). We also observed that CIT-Lasso identifies more loci in most studies (top left panel). On average, the proposed method identifies 143.6 loci whereas MAT identifies 117.1 loci (up 22.7%). This trend is more pronounced at FDR=0.2 and for phenotypes with higher polygenicity, as expected (top right panel). For example, the proposed method identifies more loci in 67.2% (89.6% at FDR=0.2) studies. The proportion increases to 81.0% (100% at FDR=0.2) for studies with more than 100 GWAS loci, where the average number of loci identified by CIT-Lasso is 349.3 vs. 279.6 for MAT (up 24.9%). We present specific results for several common polygenic traits in **Figure 7** (bottom three panels) including BMI (Yengo et al. 2018; 610 vs. 376 loci), height (Yengo et al. 2018; 714 vs. 639 loci), total cholesterol (Willer et al. 2013; 127 vs. 76 loci), bone mineral density (Kemp et al. 2017; 262 vs. 203 loci), hypertension (Zhu et al. 2019; 260 vs. 185 loci), type 2 diabetes (Xue et al. 2018; 172 vs. 94 loci).^41–45^ The results demonstrate the generalizability of the proposed method to the phenome, and that it is particularly useful for complex traits with high polygenicity.

## Materials and Methods

### Conditional independence hypothesis and conditional causal effect

In causal inference, the causal effect of *G*_*j*_ on *Y* is defined on the basis of the counterfactual (or potential) outcomes 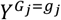 and 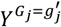, which are mutually unobservable quantities.^25,26^ 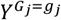 is the outcome variable that would have been observed under the genotype value 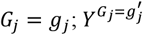 is the analogous quantity when 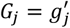. To evaluate the causal effect of the *j*-th genetic variant on *Y*, we are interested in testing the nonparametric conditional causal effect (CCE) for all joint strata *G*_−*j*_ = *g*_−*j*_:

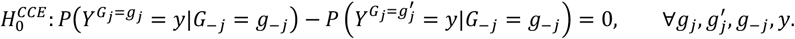

The conditional causal effect ensures that the confounding effect through other genetic variants has been adjusted for.^26^ Intuitively, the CCE evaluates the effect of *G*_*j*_ on *Y* by changing the value of *G*_*j*_ from *g*_*j*_ to 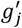, without altering any other variants. If the distribution of *Y* changes for any stratum *G*_−*j*_ = *g*_−*j*_, the *j*-th variant may be said to have a causal effect on *Y*. This is conceptually similar to a functional experiment that edits a particular sequence and then looks for any change in the phenotype (e.g., MPRA or CRISPR).

While it is natural to consider this causal inference hypothesis 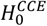, it is unclear how one should go about testing it. To address this, we consider an alternative conditional independence hypothesis which has been previously proposed to identify variants that carry non-redundant information on the phenotypes:^13,14,16,18,19^

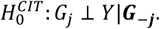

Importantly, our methodology based on model-X knockoffs offers a way to perform conditional independence test without assuming that a disease model *Y*|*X* is correctly specified (see Sections below).

We show that this hypothesis is equivalent to testing the conditional causal effect for all strata *g*_−*j*_ with *P*%*G*_−*j*_ = *g*_−*j*_) > 0, i.e., 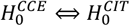, under the following conditions:

1. Unconfoundedness (conditional exchangeability): 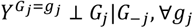.
2. Positivity: *P*(*G*_*j*_ = *g*_*j*_|*G*_−*j*_ = *g*_−*j*_) > 0 for all *g*_−*j*_ with *P*(*G*_−*j*_ = *g*_−*j*_) > 0.
3. Consistency: 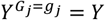 for every individual with *G*_*j*_ = *g*_*j*_, where *Y* is the observed outcome.

These three conditions are the same as the usual identification conditions in causal inference^25^, which characterize the extent to which a statistical claim (based on observable quantities) can be interpreted as a causal effect. Condition 1 assumes no unmeasured confounders beyond *G*_−*j*_. This means that the proposed test will attenuate confounding effects induced by linkage disequilibrium (the major confounding effect in genetic studies), but will not account for other unmeasured confounders such as confounding effects from genetic variants that are not genotyped or some environmental factors; Condition 2 assumes that *G*_*j*_ and *G*_−*j*_ are not deterministically linked (i.e., there must be enough variation in *G*_*j*_ for strata defined by *G*_−*j*_ = *g*_−*j*_); Condition 3 is a rule in the logic of counterfactuals as discussed by Pearl^51^ that the potential outcome under the assignment *G*_*j*_ = *g*_*j*_ is the outcome *Y* that will actually be observed in the event *G*_*j*_ = *g*_*j*_. Under the unconfoundedness assumption and the consistency condition,

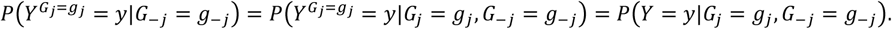

Therefore, the null hypothesis for the conditional causal effect for all joint strata *G*_−*j*_ = *g*_−*j*_,

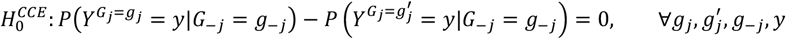

can be written in terms of observable quantities as

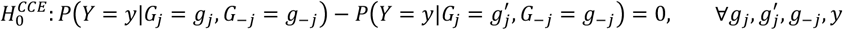

which is equivalent to

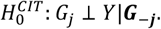

### Conditional independence test via GhostKnockoffs and genome-wide sparse regression

To test the conditional independence hypothesis on the basis of summary statistics, He et al. (2022) proposed GhostKnockoffs and showed that for a particular feature importance defined by score test statistics in marginal association models, one can directly generate the knockoff feature importance score per variant without the need to generate individual-level knockoffs for hundreds of thousands of samples.^22^ In this paper, we extend GhostKnockoffs to measures of feature importance evaluated via a genome-wide sparse regression. Compared to the original GhostKnockoffs, the sparse regression is expected to boost statistical power and improve prioritization of causal variants. We consider the setting where individual-level data ***Y*** and ***G*** are not available but summary statistics are available, including: 1. sample size of the target study; 2. marginal Z-scores from a target study; 3. correlation matrix from a reference panel. For studies where the marginal Z-scores are not readily available, they can be computed using p-values and direction of effects as *Z*_*j*_ = sign_*j*_ × Φ^−1^(*p*_*j*_/2).

Our approach builds on the results in Chen et al. (2024),^24^ and consists of four main steps: (1) generate multiple (*M*) feature importance knockoff per variant, 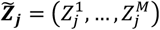 using a marginal association framework as in He et al. (2022) (2) re-calculate a new vector of feature importance measures via a genome-wide sparse regression that includes both original and knockoff variants; (3) define the test statistics (knockoff scores) by contrasting the feature importance for the original and knockoff variants; (4) implement the knockoff filter procedure to select important variants with FDR control.^24^ Here, steps (1) - are applied to each stochastically independent LD block separately (independence is here approximate). The LD blocks are defined by Berisa and Pickrell (2016).^46^

The output of the analysis contains selected genetic variants that form what we refer to as the “catching set”, analogous to the “credible set” in Bayesian fine-mapping methods. Our catching sets are genome-wide, mutually exclusive, and they represent conditionally independent effects on the outcome of interest. We have developed a computational pipeline to enable scalable implementation of the proposed method. Empirically, it only requires 860.5 seconds to meta-analyze ten AD genetic studies with a single Intel Xeon E5-2640 CPU (2.4GHz) with 24GB of requested RAM.

#### Step 1: Generate knockoff Z-scores

Assuming we generate *M* knockoff copies, He et al. (2022)^22^ showed that one can generate knockoff feature importance measure according to a Z-score test by

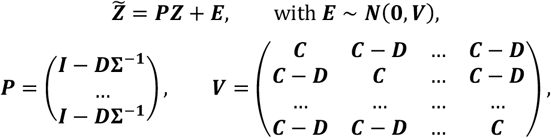

where 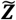 is a *pM*-dimensional vector; ***I*** is a *p* × *p* identity matrix; **Σ** is the correlation matrix of ***G***_***i***_ that characterizes the linkage disequilibrium; ***C*** = 2***D*** − ***D*Σ**^−**1**^***D***; ***D*** = diag(*s*_1_, …, *s*_*p*_) is a diagonal matrix that can be chosen to maximize the “difference” between original variables and their knockoffs, to increase the power of discovering importance variables.^14,22^

Unfortunately, when variables are strongly correlated, the requirement that ***V*** is a symmetric, positive definite matrix results in very limited range available for *s*_1_, …, *s*_*p*,_, with negative impact on power. This is a reflection of the fact that while conditional independence hypotheses aim to resolve the contribution of each individual variant, avoiding signals due to LD, this can result in loss of power in presence of tightly linked variants, as it becomes impossible to distinguish true causal genetic variants from others highly correlated with them on the basis of a limited sample size. To address this issue, it is useful to identify groups of highly correlated variants, and modify the hypotheses to test whether variants in a group have an effect on the response, conditional on all variants outside the group.^13,18^ In practice, groups are defined by applying average linkage hierarchical clustering with correlation coefficient cutoff 0.5.

This modification of the inferential target results in a different requirement on the joint distribution of the original and knockoff variables, referred to as “group knockoffs”. Here we follow the Chu et al. (2023)^23^. Unlike the single-variant knockoff construction above where ***D*** is a diagonal matrix, group knockoffs consider that ***D*** is a block diagonal matrix where each block corresponds to a predefined group of tightly linked variants. Intuitively, we aim to find “large” ***D*** such that the knockoffs are sufficiently different from the original data to achieve high power (note that the fact that ***D*** is block diagonal, rather than diagonal gives us more freedom). Among various possibilities considered in the literature, maximizing the entropy exhibits promising empirical power. It translates to solving the following convex optimization problem:

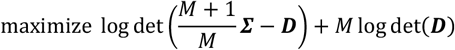

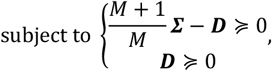

where the objective function is the entropy of the joint distribution and the constraint ensures that ***V*** is a positive definite matrix. To solve this problem, we apply the coordinate descent algorithm for maximum entropy group knockoffs and accelerate its convergence by exploiting the conditional independence among groups, following Chu et al. (2023).^23^

Finally, to generate ***E*** ∼ ***N***(**0, *V***), one classically uses a Cholesky decomposition of ***V***, which can be computationally intensive since ***V*** is an *Mp* × *Mp* matrix. Here, we utilize the structure of ***V*** and propose an efficient sampling method as proposed in Yu et al. (2024) and described as follows.^47^

1. Compute the Cholesky factorization 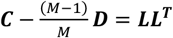.
2. Generate ***e***_1_ ∼ ***N***(**0, *IP***); let 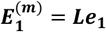.
3. Generate 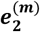 from *N*(0, ***D***); Compute 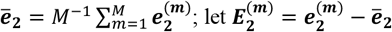.
4. Calculate ***E*** = ***E***_**1**_ + ***E***_**2**_.

As a result, the computational complexity of sampling ***E*** is reduced from *O*(*M*^3^*p*^3^) to *O*(*p*^3^).

#### Step 2: Calculate the importance of original and knockoff features using ultrahigh-dimensional sparse regression

We use an ultrahigh-dimensional sparse regression as a tool to analyze data and to quantify feature importance. It is worth noting that knockoff inference holds without assuming that this model is correctly specified. We adopt the objective function proposed by Chen et al. (2024) that minimizes:

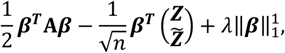

where 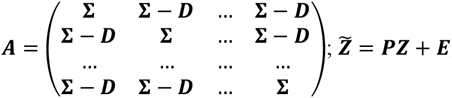 with ***E*** ∼ ***N***(**0, *V***).^24^ To address the issue that ***A*** can be nearly singular and the solution can be numerically unstable, in practice we work with ***A*** + 0.01***I***_***A***_ where ***I***_***A***_ is an (*M* + 1)*p* × (*M* + 1)*p* identity matrix. The shrinkage version is numerically equivalent to an elastic-net problem:

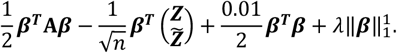

Once 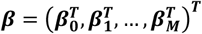 is estimated, we can then calculate the feature importance score as the magnitude of effect size: 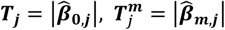. This optimization approximates the penalized regression using individual level data. The resulting feature importance score is substantially more powerful than that used in the original version of GhostKnockoffs developed by He et al. (2022) on the basis of marginal Z-scores.^22^

Solving the optimization problem defined by this objective function is a non-trivial task when the number of genetic variants is very large. We adopt a fast and scalable batch screening iterative lasso (BASIL)^48^ algorithm to solve 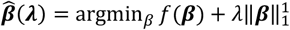, where 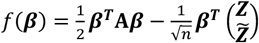. Compared to the previous BASIL algorithm based on individual level data, our BASIL algorithm requires only summary statistics and leverages the structure of the matrix ***A*** for substantially improved computational efficiency. The algorithm is described as follows.

**Initialization**: Let 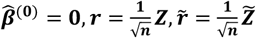.

1. Compute 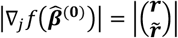 and 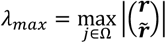.
2. Create a sequence of λ, Λ = {λ_*max*_ = λ_0_ > … > λ_*K*_} equally spaced in log-scale. Let Λ^(1)^ = {λ_0_, …, λ_*B*−1_} be the first *B* values.
3. Find *M* variants with the largest 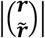. Denote the index set as *S*^(1)^.

**Iteration:** for the *k*-th iteration, *k* ≥ 1,

1. Solve the optimization problem for Λ^(*k*)^, *S*^(*k*)^ to get 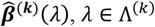.
2. Compute 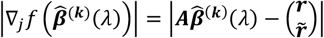, for each λ ∈ Λ^(*k*)^.
3. Find smallest λ ∈ Λ^(*k*)^, denoted as λ_*k,KKT*_, such that 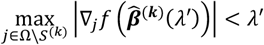 for all λ ≤ λ′ ∈ Λ^(*k*)^.
4. Let *j*^(*k*)^ denote the index at which λ_*k,KKT*_ appears in the original sequence Λ. Create a new sequence 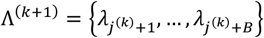.
5. Find the Δ*M*_*k*_ variables in Ω\*S*^(*k*)^ with largest 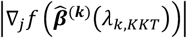. Add the Δ*M*_*k*_ variables to *S*^(*k*)^. Denote the updated index set as *S*^(*k*+1)^ for the next iteration.
6. Repeat the iterations until λ_*k,KKT*_ reaches the smallest value λ_*K*_.

To select the tuning parameter in absence of individual level data, we adopted the method proposed by Chen et al. (2024):

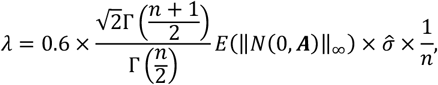

where 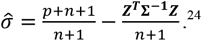.^24^ For data from genetic association studies, we observed that σ is often estimated to be very close to 1; the extreme value *E*(‖*N*(0, ***A***)‖_∞_) can be well approximated by *E* (‖*N*%0, ***I***_(*M*+1)*p*_)‖_∞_) due to the approximately block-diagonal LD structure, where ***I***_(*M*+1)*p*_ is an (*M* + 1)*p* × (*M* + 1)*p* identity matrix; 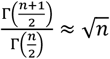 when *n* is sufficiently large. Therefore, we used the approximation below:

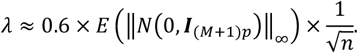

We calculated *E*(‖*N*%0, ***I***_(*M*+1)*p*_)‖_∞_) by Monte Carlo simulation with 10 replicates. Notably, this approximation can be calculated prior to data analysis, enabling efficient and robust parallel analysis of approximately independent LD blocks with the same tuning parameter. While this paper focuses on a Lasso-type model implementation, it is worth noting that the proposed framework can also flexibly leverage the SuSiE-type model and utilize its predictions as a test statistic in lieu of the Lasso to simultaneously perform discovery and prioritization of causal variants.

#### Step 3: Calculate knockoff scores (test statistics)

After the feature importance are calculated, we compute the statistics

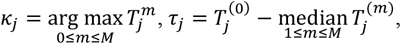

where *m* indicates the *m*-th knockoff copy and 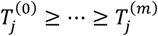 are the order statistics of 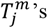 where 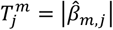 as described in Step 2. For the *j*-th variant, *κ*_*j*_ denotes the index of the original (denoted as 0) or the knockoff feature that has the largest importance score; *τ*_*j*_ denotes the difference between the largest importance score and the median of the remaining importance scores. Here, knockoff scores *κ* and *τ* obey a property similar to the “flip-sign” property in the case of a single knockoff copy.^16,49^ In the multiple knockoff scenario, *κ*_*j*_ plays the role of the sign, and *τ*_*j*_ quantifies the magnitude of the contrast. Subsequently, we define a test statistic to quantify the magnitude of effect on the outcome as

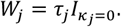

Variants with *W*_*j*_ > *t*_*α*_ are selected, where *t*_*α*_ is the threshold calculated by the knockoff filter at target FDR= *α*.

#### Step 4: Apply modified variant-level group knockoffs filter

To provide variant level interpretation given group knockoffs, we adopt a modified knockoff filter proposed by Gu et al. (2024) to test for

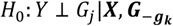

where *g*_*k*_ represents the group where the *j*-th variant resides; 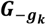 are the genotypes for all variants across the genome except variants in *g*_*k*_.^50^ Intuitively, this hypothesis corresponds to conditional independence tests between groups and marginal association tests within each group (so the variants within a group are not conditioned on each other). The modified knockoff filter at target FDR= *α* is defined as

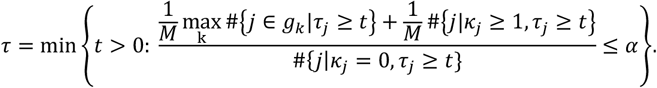

In addition, we define the q-value for a variant with statistics *κ* = 0 and *τ* as

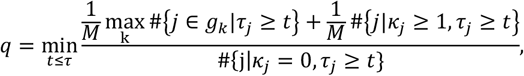

where 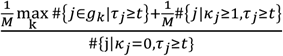 is an estimate of the proportion of false discoveries if we are to select all variants with feature statistics *κ*_*j*_ = 0, *τ*_*j*_ ≥ *t*. For variants with *κ* ≠ 0, we define *q* =1 and they will never be selected. Selecting variants with *W* > *τ*, where *τ* is calculated at target FDR= *α*, is equivalent to selecting variants with *q* ≤ *α*. The proof of this variant-level group knockoffs filter is provided in Supplementary Materials. Compared to the group knockoff filter that selects the entire group that contains the causal variants as the catching set, this variant-level filter highlights variants within each group that exhibit larger feature importance. In practice, we found that this significantly reduces the size and improves the purity of the catching set when it is paired with our proposed ultrahigh-dimensional sparse regression which additionally introduces sparsity to the feature importance scores within each group.

### Connections with other causal inference methods

Our task differs from conventional applications of causal inference on treatment effects. Although many standardization and inverse probability weighting methods have been developed to adjust for potential bias in observational studies, most existing approaches focus on inference on the effects of binary treatments in a one-by-one (i.e., marginal) manner.^26,51–59^ Such approaches are not applicable to large-scale genetic studies where the goal is to simultaneously infer the causal effect of a large number of genetic variants on the outcome of interest – indeed, extensions of causal inference methodology to high-dimensional biology applications is a limited but growing area.^60–63^ This motivates our proposed method based on CCE and knockoffs to mimic a functional experiment that edits a particular sequence and looks for any change in the phenotype. In this section, we clarify the connections between our method and other existing causal inference methods.

#### Unconditional causal effect (UCE)

One popular measure of causal effect in epidemiologic studies is the UCE.^26^ The corresponding hypothesis is defined as

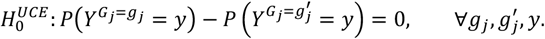

If a variant obeys the null hypothesis 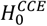, it also obeys the null 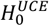 because

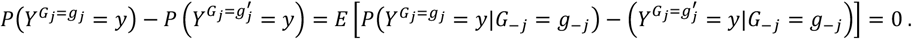

The implication is naturally one-sided: the null hypothesis 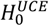 does not imply the null 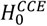. That is, the proposed method may successfully identify a genetic variant exhibiting a causal effect in one stratum, but the said variant may not have an *unconditional* causal effect (i.e., when marginalizing across strata).

#### Average causal effect (ACE).^26^

An analogous derivation can be applied to the average causal effect 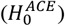, defined as

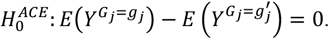

If the underlying model is linear, *Y* = ∑_*j*_ *β*_*j*_*G*_*j*_ + *ε*, testing conditional independence is equivalent to testing the unconditional causal effect and testing the average causal effect under the same three identification conditions, i.e.,

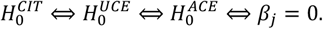

#### Mendelian randomization

Our task of finding causal variants substantially differs from that of Mendelian randomization, another popular causal inference method for genetic data.^64–66^ Unlike MR, which uses genetic variation as an instrumental variable to investigate the causal effect of a modifiable exposure (e.g., demographic features, gene expressions, proteins) on the outcome of interest subject to unmeasured confounders, our target directly lies on the causal pathway between genetic variation and the outcome of interest under the unconfoundedness assumption.

### Meta-analysis of potentially overlapping studies

Our proposed method takes marginal Z-scores from a target study as input. Here we describe a simple and effective method to calculate meta-analysis Z-scores by aggregating Z-scores from possibly overlapping studies. The calculated meta-analysis Z-score directly serves as the input of the proposed knockoff inference, which allows the integration of multiple studies to the maximum extent feasible. We note that this significantly simplifies the meta-analysis pipeline proposed by He et al. (2022) because one no longer needs to perform knockoff inference for each study separately.^22^

Let ***Z***_***k***_ be the Z-scores from the *k*-th study and *n*_*k*_ be the sample size of the *k*-th study. We define meta-analysis Z-score as:

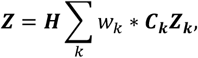

where the optimal weights *w*_*k*_ are given by solving

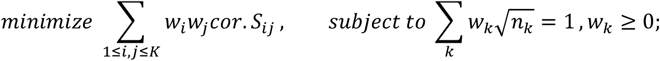

***C***_***k***_ = *diag*(*c*_*k*1_, …, *c*_*kp*_) is a diagonal matrix where *c*_*kj*_ = 1 if *Z*_*kj*_ is observed and *c*_*kj*_ = 0 otherwise; ***H*** = *diag*(*h*_1_, …, *h*_*p*_) where 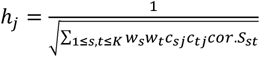. We calculate the study correlation matrix

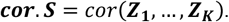

In practice, we use variants with |Z-score|≤1.96 to calculate ***cor*.*S*** to remove the correlation due to polygenic effects. Similar methods have been proposed by Lin and Sullivan (2009) and implemented in METAL (https://genome.sph.umich.edu/wiki/METAL_Documentation).^67,68^

## Discussion

The described method for searching for causal variants, CIT-Lasso, combines ideas from causal inference, model-X knockoffs and ultrahigh-dimensional sparse regression. By applying principles of causal inference, we designed a high-dimensional hypothesis testing problem that targets the same causal estimand as experimentally editing a particular genomic sequence. We subsequently transformed the initial causal inference hypothesis into a conditional independence hypothesis, which can be rigorously tested by the model-X knockoffs framework. We facilitated this test by fitting an innovative ultrahigh-dimensional sparse regression, which enables the simultaneous modeling of all genomic variants genotyped in the human genome. This results in a data-driven method that learns far more effectively and accurately from genetic data compared to the conventional marginal association models that analyzes one genetic variant at a time, and then proceeds to a second stage fine-mapping procedure. We implemented our approach in a computationally efficient open-source software allowing a genome-wide analysis to be performed in less than 15 minutes on a single CPU. Crucially, we require only summary statistics: our approach can be flexibly applied on top of the standard GWAS pipeline without changing any of the processing steps, resulting in the discovery of additional loci and enhanced localization of causal variants.

The superior power compared to conventional marginal association tests can be attributed to several factors including: 1. the use of FDR control as opposed to the stringent genome-wide significance level for marginal association tests (5 × 10^−8^); and 2. the use of an ultrahigh-dimensional sparse regression that jointly models all genetic variants, thereby making inference more efficient. Both are especially helpful for complex traits with high polygenicity. In particular, ultrahigh-dimensional sparse regression has been employed recently in genetic research to establish polygenic risk scores that predict risk of complex diseases.^48^ It is well known that inclusion of genetic variants with relatively weaker associations that do not meet the stringent genome-wide significance level can significantly improve a model’s prediction accuracy. This aligns with our observations that the inference derived from the proposed ultrahigh-dimensional sparse regression exhibits improved power relative to marginal association tests. However, it is difficult to conduct hypothesis testing in an ultrahigh-dimensional sparse regression^69^ and it is challenging to control FDR for interpretable discoveries in GWAS.^27^ The integration with model-X knockoffs makes rigorous inference possible. This is a salient advantage of our proposed methodology compared to the original GhostKnockoffs method, which conducts similar conditional independence tests with marginal association test statistics.^22^

Existing fine-mapping methods calculate relative likelihoods of causality using the PIP (posterior inclusion probability).^10^ PIP is a ranking measure that indicates how likely a predictor is to be included in a model, lacking a systematic approach to defining a cutoff for determining causal variants.^10,11,70^ By contrast, the proposed method is an end-to-end method that simultaneously performs discovery and prioritization of causal variants. While it makes more discoveries, it is also superior to state-of-the-art second-stage fine-mapping methods in pinpointing the causal variants, and it automatically determines a cutoff that controls the FDR. Moreover, the current implementation of the ultrahigh-dimensional sparse regression can be replaced by any existing and future fine-mapping models and provide an inferential cutoff for PIP, as illustrated in the simulation studies (CIT-SuSiE), leveraging thier possibly enhanced performance in localizing causal variants.

Another appealing feature of the proposed method is that the identified variants per locus are grouped into distinct sets, with conditionally independent effects on the phenotype of interest. This allows the identification of independent causal effects underlying association signals with FDR control. As a proof of concept, in **Supplementary Figure 6**, we present the number of identified variants, number of identified conditional independent effects (defined as groups of identified variants) and number of mapped genes via cS2G for each locus. We observed that as the number of conditionally independent effects increases, the number of mapped genes correspondingly increases (correlation coefficient = 0.81). This shows that the different conditionally independent effects tend to be mapped to different genes that they potentially regulate. In **Supplementary Figure 7**, we present some concrete examples for *BIN1, PTK2B* (*CLU*) and *HLA* complex. For *BIN1*, the proposed method identified three conditionally independent effects, which are mapped to *PROC* (2 variants), *BIN1* (1 variant) and *ERCC3/BIN1* (3 variants) respectively; for *PTK2B*, the proposed method identified two conditionally independent effects, which are mapped to *PTK2B* (2 variants) and *CLU* (1 variant); for HLA complex, the proposed method identified five conditionally independent effects that can be mapped via cS2G to five different genes *HLA-DRB5* (1 variant), *HLA-DQA2* (1 variant), *SLC44A4* (1 variant), *AIF1* (1 variant), *POU5F1* (1 variant). The findings are consistent with previous studies showing that genetic association signals can manifest through multiple causal variants with conditionally independent effects on the phenotype of interest via regulating different genes.^12^

A limitation of our primary analysis is that we only considered directly genotyped variants and hence the causal variants can be missing from the analysis. While imputation has been shown to improve the statistical power of marginal association models, the situation changes when our goal is to pinpoint the causal variants. The imputed genotypes do not carry information on the phenotype in addition to the one in the directly genotyped variants, on the basis of which they have been imputed. This means that all the conditional independence hypotheses corresponding to the imputed variants are null.^13^ Using a different perspective, formal causal inference is solely viable for variants that are directly genotype. Indeed, imputed variants violate the positivity condition (see **Methods**), since they always operate as a function of other variants and remain independent of the phenotype of interest when conditioned on the variants used for imputation. Nevertheless, with appropriate qualifications, it is possible to introduce imputed variants in the study and we additionally performed a comparison between the analyses with directly genotyped variants (our primary analysis) and the analyses with all imputed variants. Although it is commonly assumed that imputation can facilitate fine-mapping the causal variant, we found that imputation does not improve fine-mapping in the nine examples considered here for SuSiE and CIT-Lasso. For the three examples where the causal variants are directly genotyped, SuSiE and CIT-Lasso can no longer identify the causal variants (**Supplementary Figure 3**). For the six examples where the causal variants are not directly genotyped, SuSiE and CIT-Lasso also fail to pinpoint the causal variants (causal variants are not included in any credible/catching sets) except for one example rs636317 (**Supplementary Figure 4**). Our results emphasize the significance of whole-genome sequencing data in pinpointing causal variants rather than simply imputing. As whole genome sequencing data becomes increasingly available, we hope that the proposed method and computational approaches that apply its techniques will become essential tools for identifying causal biological pathways. This, in turn, should expedite the creation of new therapeutic interventions.

## Data Availability

Summary statistics for Alzheimer’s disease: 1. The genome-wide survival association study performed by Huang et al. (2017) (https://www.niagads.org/datasets; NIAGADS ID: NG00058); 2. The genome-wide meta-analysis by Jansen et al. (2019) (https://ctg.cncr.nl/software/summary_statistics); 3. The genome-wide meta-analysis by Kunkle et al. (2019) (NIAGADS ID: NG00075); 4. The genome-wide meta-analysis by Schwartzentruber et al. (2021) (https://www.ebi.ac.uk/gwas/; GWAS catalog ID: GCST90012877); 5. In-house genome-wide association study imputed using the TOPMed reference panels (see Supplementary Table 3); 6-7. Two whole-exome sequencing analyses of data from ADSP by Bis et al. (2020) (NIAGADS ID: NG00065), and Le Guen et al. (2021) (NIAGADS ID: NG000112); 8. In-house whole-exome sequencing analysis of ADSP (NIAGADS ID: NG00067.v5); 9. In-house whole-genome sequencing analysis of ADSP (NIAGADS ID: NG00067.v5); 10. The genome-wide meta-analysis by Bellenguez et al. (2022) (GWAS Catalog ID: GCST90027158).

Summary statistics of the 400 GWAS: https://github.com/mikegloudemans/gwas-download.

Pan-UKBB reference panel: https://pan.ukbb.broadinstitute.org/.

The results of our analysis can be downloaded at https://github.com/biona001/ghostknockoff-gwas-reproducibility.

## Code Availability

Our overall method is implemented in the software GhostKnockoffGWAS, which is available at https://github.com/biona001/GhostKnockoffGWAS. The high-dimensional Lasso regression for summary statistics data is implemented in an independent software ghostbasil, which is itself a standalone R package, available at https://github.com/JamesYang007/ghostbasil. Pre-computed knockoff statistics from the European panel of Pan-UKB are freely available for download. Finally, scripts to reproduce the results in this paper can be accessed at https://github.com/biona001/ghostknockoff-gwas-reproducibility.

## Acknowledgements

This research was additionally supported by NIH/NIA award AG066206 (ZH), AG066515 (ZH), AG075238 (MEB), EB001988-21 (TH, JY), and by the Simons Foundation under award 814641 (ZC). We gratefully acknowledge the studies which provided summary statistics.

## Author Contributions

Z.H., C.S. and E.C. developed the concepts for the manuscript. Z.H., B.C., J.Y. and J.G. proposed the methods. C.S., E.C., I.I.-L and T.H. supervised the method development. Z.H. designed the analyses and applications. Z.C., T.M., L.L., N.H. and M.M. significantly improved the statistical methods. I.I.-L and H.T. significantly improved the design of the analyses and applications. Z.H., B.C., M.E.B. and Q.X. conducted the analyses. Y.L. helped with interpretation of AD genetics. Z.H. prepared the manuscript and all authors contributed to editing the paper.

## Competing interests

The Authors declare no competing interests.

## Notes

### Competing Interest Statement

The authors have declared no competing interest.

### Summary of Updates

Simulation results got updated. Figures related to simulation and method comparisons are revised

