## Supplementary Materials for "Beyond guilty by association at scale: searching for causal variants on the basis of genome-wide summary statistics"

**Supplementary Figure 1: Meta-analysis of Alzheimer's disease genetic studies via marginal association tests.** We present the Manhattan plot from conventional marginal association tests. For the marginal association tests (top panel), the dotted line presents the conventional genome-wide p-value threshold of $5 \times 10^{-8}$. P-values and W statistics are truncated at $10^{-50}$ for better visualization. The results are based on p-values calculated via the proposed meta-analysis strategy with optimal weights combining the ten studies. Variant density is shown at the bottom of plot (number of variants per 1Mb).

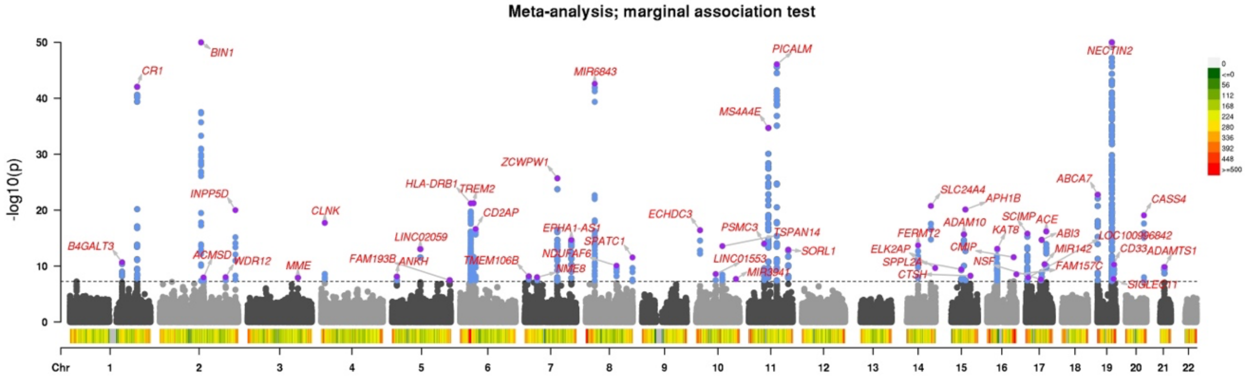

**Supplementary Figure 2: Conditional independence test via CIT-Lasso consistently identifies more loci and enables prioritization of putative causal variants.** We applied the proposed method to three major meta-analyses of AD genetic studies from 2019 to 2022. Finally, we performed a meta-analysis of 10 AD genetic studies from 2017-2022. We report the number of identified loci and the number of identified proxy variants per loci.

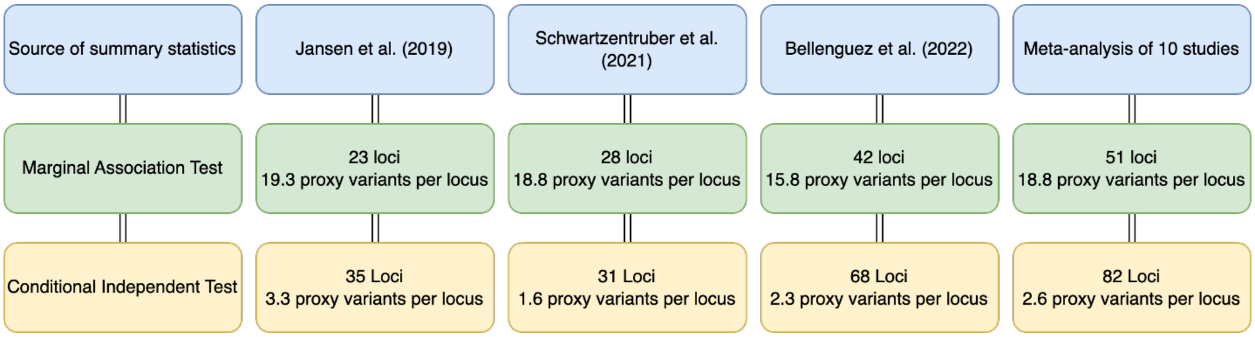

**Supplementary Figure 3. Fine-mapping with directly genotyped data vs. imputed data when the MPRA and CRISPR validated variants are typed.** We compare marginal association test (MAT), MAT followed by SuSiE fine-mapping, and CIT-lasso. Different colors represent different catching/credible sets (for CIT-Lasso, they represent independent conditional causal effects). The legend presents the genes that each catching/credible set potentially regulates, mapped by the cS2G method. The red dotted lines represent the p-value threshold of $5 \times 10^{-8}$ and FDR threshold of 0.10 for MAT and CIT-lasso respectively.

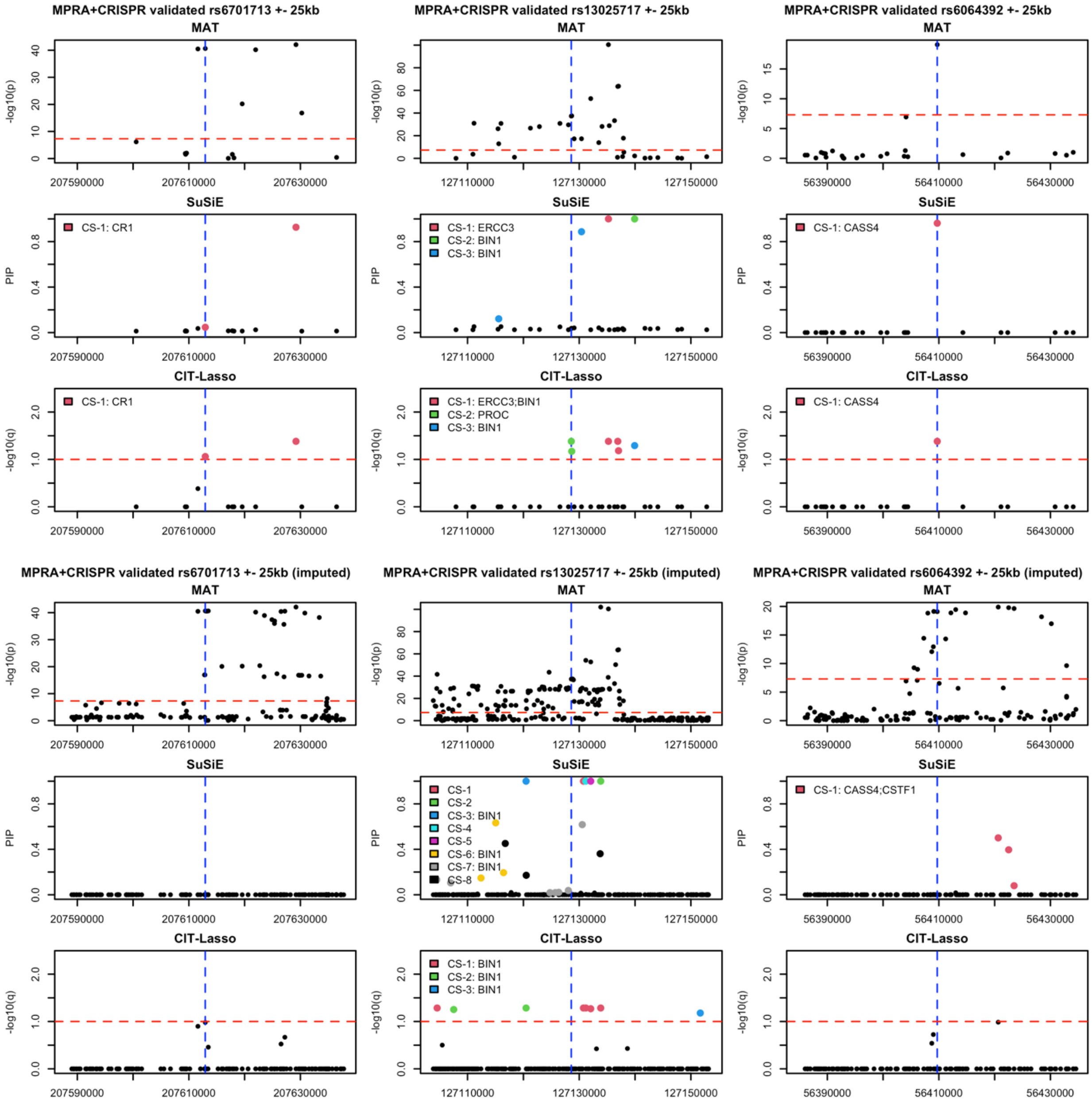

**Supplementary Figure 4. Fine-mapping with directly genotyped data vs. imputed data when the MPRA and CRISPR validated variants are not genotyped**. We compare marginal association test (MAT), MAT followed by SuSiE fine-mapping, and CIT-lasso. Different colors represent different catching/credible sets (for CIT-Lasso, they represent independent conditional causal effects). The legend presents the genes that each catching/credible set potentially regulates, mapped by the cS2G method. The red dotted lines represent the p-value threshold of $5 \times 10^{-8}$ and FDR threshold of 0.10 for MAT and CIT-lasso respectively.

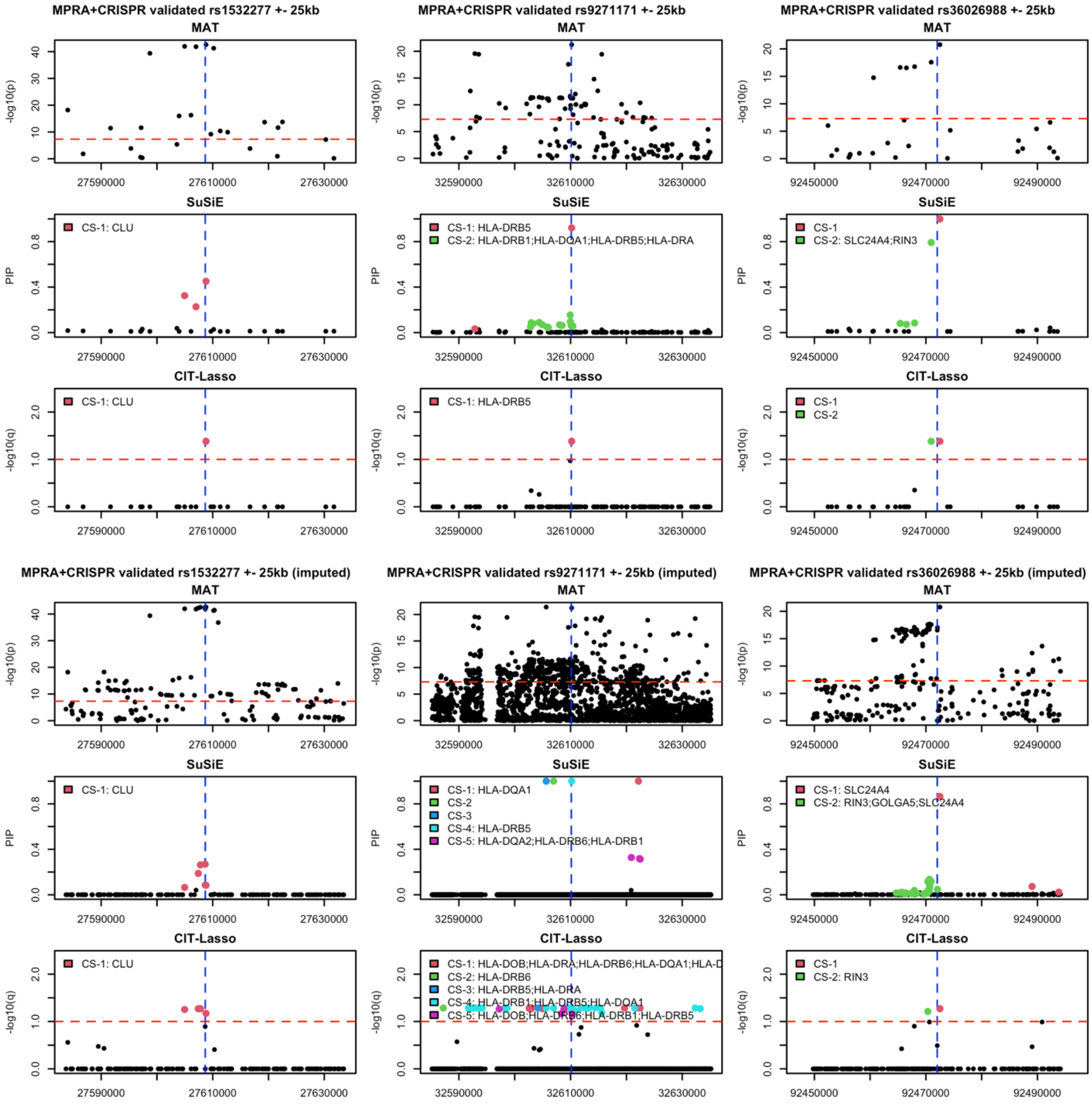

**Supplementary Figure 4 (continue). Fine-mapping with directly genotyped data vs. imputed data when the MPRA and CRISPR validated variants are not genotyped**. We compare marginal association test (MAT), MAT followed by SuSiE fine-mapping, and CIT-lasso. Different colors represent different catching/credible sets (for CIT-Lasso, they represent independent conditional causal effects). The legend presents the genes that each catching/credible set potentially regulates, mapped by the cS2G method. The red dotted lines represent the p-value threshold of $5 \times 10^{-8}$ and FDR threshold of 0.10 for MAT and CIT-lasso respectively.

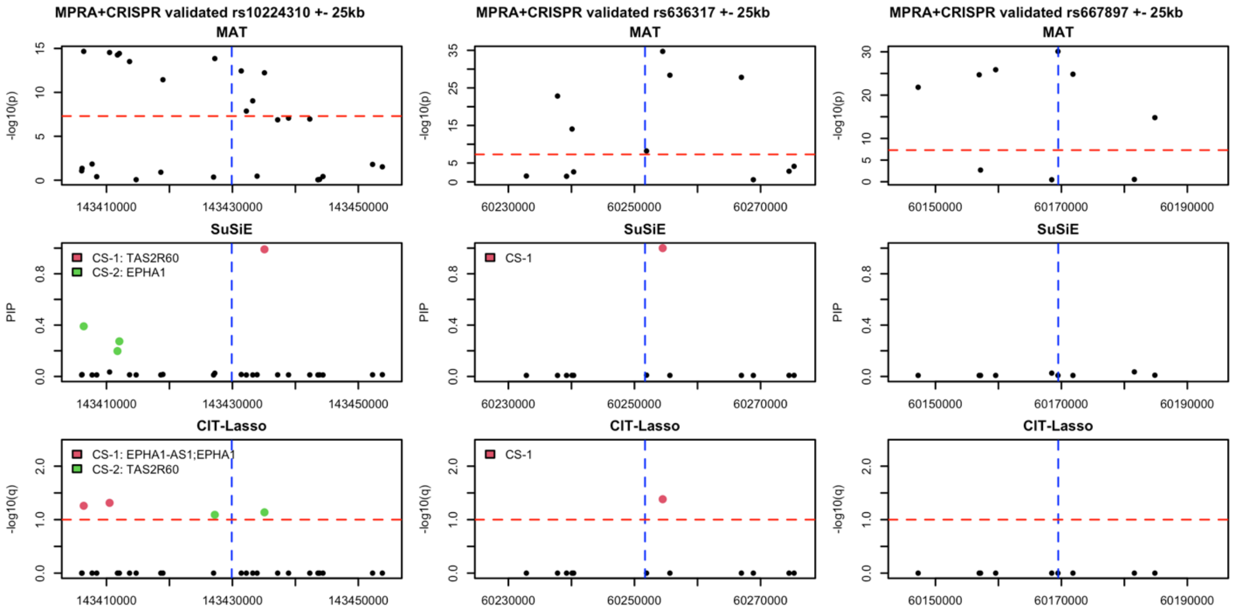


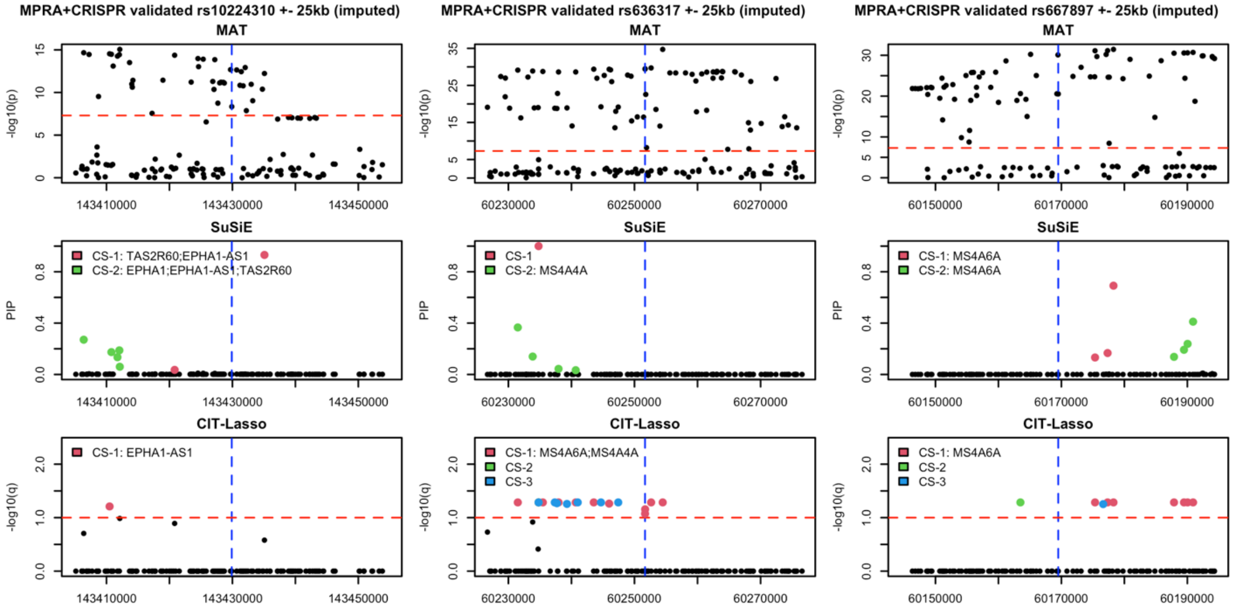

**Supplementary Figure 5. Additional variants identified by CIT-Lasso ($p \geq 5 \times 10^{-8}$) have concordant z-scores and directions of effects across 10 studies.** Left panel: we compute the spearman correlation of Z-scores between each pair of studies. Right panel: for each identified variant, we compute and present the proportion of studies with same direction of effect as in the meta-analysis.

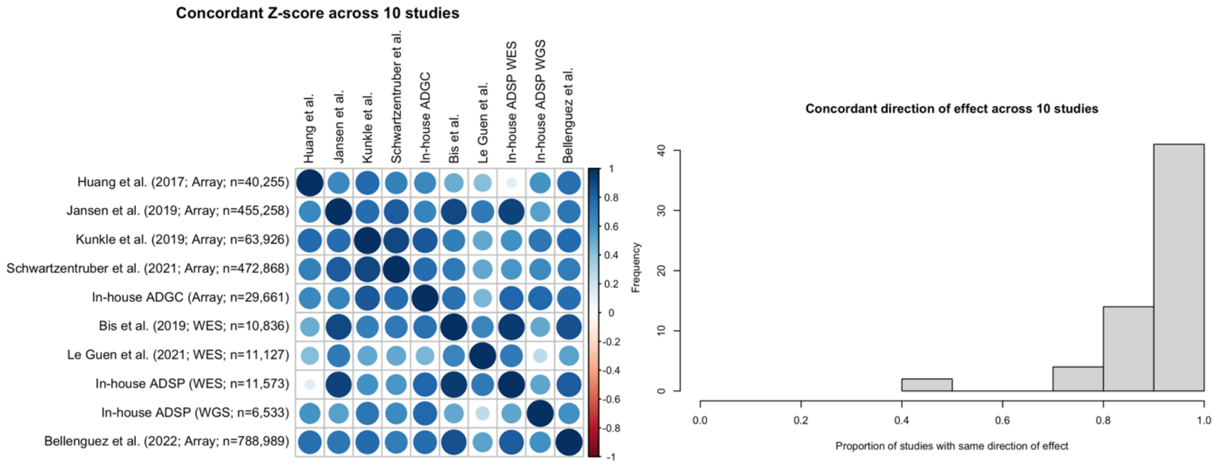


**Supplementary Figure 6. CIT-Lasso identifies conditionally independent effects that regulate distinct genes.** Left panel: Histogram of the number of conditionally independent effects (CIE) per locus. Right panel: for identified variants that can be mapped to cS2G genes, we present the number of identified variants per locus, number of identified conditional independent effects per locus and the number of corresponding cS2G genes. Each bar presents a locus. The loci are ordered by the number of identified variants. APOE locus is excluded for better visualization. The correlation between number of identified conditional independent effects and number of corresponding cS2G genes is 0.81.

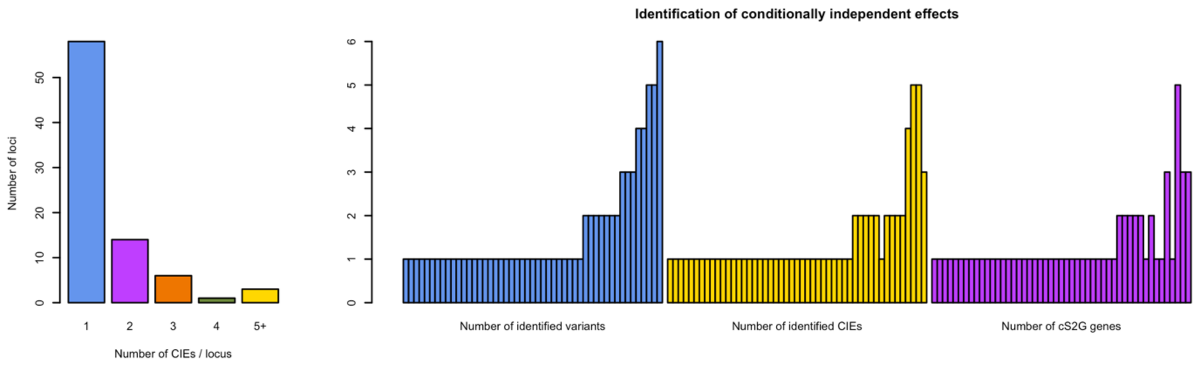

**Supplementary Figure 7. Additional fine-mapping examples for *BIN1*, *PTK2B* and *HLA-DRB1*.** We compare marginal association test (MAT), MAT followed by SuSiE fine-mapping, and CIT-lasso. Different colors represent different catching/credible sets (for CIT-Lasso, they represent independent conditional causal effects). The legend presents the genes that each catching/credible set potentially regulates, mapped by the cS2G method. The red dotted lines represent the p-value threshold of $5 \times 10^{-8}$ and FDR threshold of 0.10 for MAT and CIT-lasso respectively.

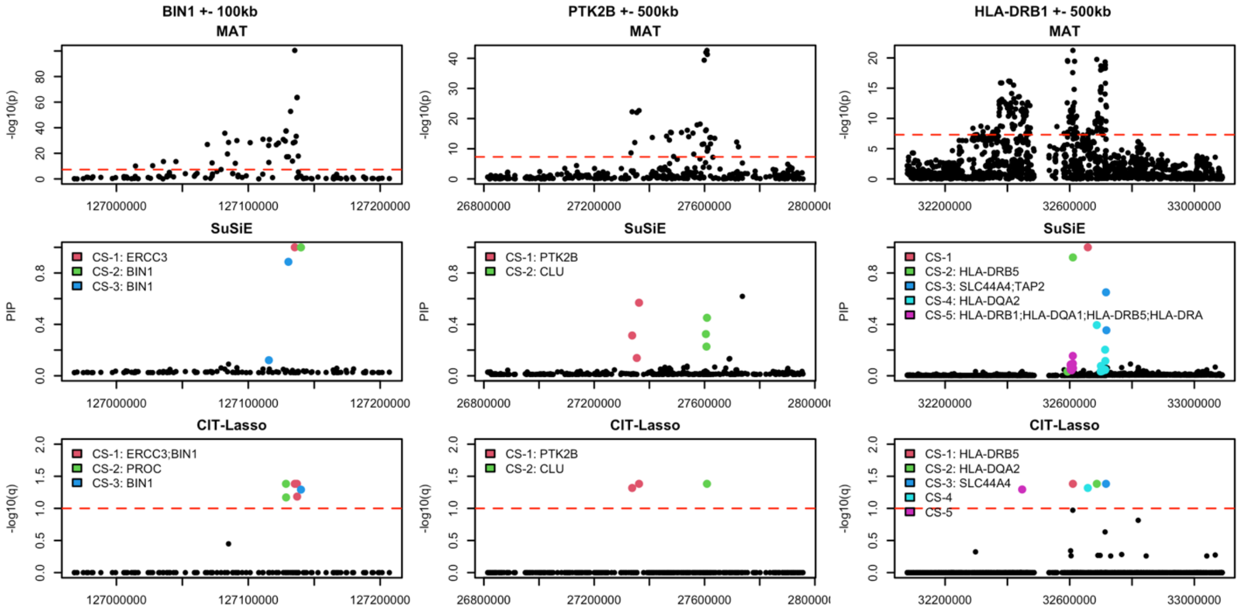

**Proof of the variant-level group knockoffs filter**

To provide variant level interpretation given group knockoffs, we consider the hypotheses:

$$H_0: Y \perp G_j | \boldsymbol{G}_{-g_k}$$

where $g_k$ represents the group in which the $j$-th variant resides; $\boldsymbol{G}_{-g_k}$ are the genotypes for all variants across the genome except variants in $g_k$. For notational simplicity, we prove the case when $M = 1$ where $\kappa_j \in \{0,1\}$. The extension to $M > 1$ follows the analogous argument for multiple simultaneous knockoffs.

For a target FDR of $\alpha$, we aim to find $t_\alpha$ such that

$$E\{FDP(t_\alpha)\} = E\left\{\frac{\#\{j \in H_0 | \kappa_j = 1, \tau_j \geq t_\alpha\}}{1 \vee \#\{j | \kappa_j = 0, \tau_j \geq t_\alpha\}}\right\} \leq \alpha$$

The existing proof of FDR control of the knockoff filter does not apply here because we are using group knockoffs. We consider a decomposition of the FDP corresponding to a partition of the variants such that the conventional knockoff filter can be applied to each partition. To achieve this, we assign all variants to an $L \times K$ matrix, where each column corresponds to one group and each row contains only one variable from each group. For each column, we rank variable within a group by $\tau_j$ and assign them in descending order. This way $\#\{j \in R_l, j \in H_0 | \kappa_j = 0, \tau_j \geq t_\alpha\} = 0$ for any $l > \phi(t_\alpha) = \max_k \#\{j \in g_k | \tau_j \geq t_\alpha\}$, where $j \in R_l$ means the $j$-th variant is in the $l$-th row. For the feature statistic defined by $T_j^m = |\hat{\beta}_{m,j}|$ where $\hat{\beta}_{m,j}$ is the coefficient estimate from lasso regression, we have the following lemma.

**Lemma**. (1) Suppose $Y \perp \boldsymbol{G}_{g_k} | \boldsymbol{G}_{-g_k}$, then $\kappa_j \sim unif\{0,1\}$ conditional on $\tau_j$ for all $j \in g_k$; (2) Suppose $Y \perp \boldsymbol{G}_{g_k} | \boldsymbol{G}_{-g_k}$ and $Y \perp \boldsymbol{G}_{g_{k'}} | \boldsymbol{G}_{-g_{k'}}$, $k \neq k'$, then $\kappa_j \perp \kappa_{j'} | \tau_1, \ldots, \tau_p$ for any pairs $j \in g_k$ and $j' \in g_{k'}$, $k \neq k'$.

Proof to (1): By the definition of group knockoffs,

$$y \perp (\widetilde{\boldsymbol{G}}_{g_k}, \widetilde{\boldsymbol{G}}_{-g_k}) | \boldsymbol{G}_{g_k}, \boldsymbol{G}_{-g_k}.$$

Since $y \perp \boldsymbol{G}_{g_k} | \boldsymbol{G}_{-g_k}$,

$$y \perp (\boldsymbol{G}_{g_k}, \widetilde{\boldsymbol{G}}_{g_k}, \widetilde{\boldsymbol{G}}_{-g_k}) | \boldsymbol{G}_{-g_k}$$

$$\Rightarrow y \perp (\boldsymbol{G}_{g_k}, \widetilde{\boldsymbol{G}}_{g_k}) | \boldsymbol{G}_{-g_k}, \widetilde{\boldsymbol{G}}_{-g_k}$$

By the definition of group knockoffs,

$$(\boldsymbol{G}_{g_k}, \widetilde{\boldsymbol{G}}_{g_k}) | \boldsymbol{G}_{-g_k}, \widetilde{\boldsymbol{G}}_{-g_k} \sim (\widetilde{\boldsymbol{G}}_{g_k}, \boldsymbol{G}_{g_k}) | \boldsymbol{G}_{-g_k}, \widetilde{\boldsymbol{G}}_{-g_k}$$

where $\sim$ denotes equality in distribution. Therefore,

$$(G_j, \tilde{G}_j, \boldsymbol{G}_{g_k \backslash j}, \widetilde{\boldsymbol{G}}_{g_k \backslash j}, \boldsymbol{G}_{-g_k}, \widetilde{\boldsymbol{G}}_{-g_k}, y) \sim (\tilde{G}_j, G_j, \widetilde{\boldsymbol{G}}_{g_k \backslash j}, \boldsymbol{G}_{g_k \backslash j}, \boldsymbol{G}_{-g_k}, \widetilde{\boldsymbol{G}}_{-g_k}, y)$$

Subsequently,

$$T_j^0 = |\hat{\beta}_{0,j}| \sim |\hat{\beta}_{1,j}| = T_j^1 | \tau_1, \ldots \tau_p$$

$$\Rightarrow \kappa_j \sim unif\{0,1\} | \tau_1, \ldots \tau_p$$

for all $j \in g_k$.

Proof of (2): It suffices to prove

$$\left(T_j^0, T_j^1, T_{j'}^0, T_{j'}^1\right) \Big| \tau_1, \ldots \tau_p \sim \left(T_j^1, T_j^0, T_{j'}^0, T_{j'}^1\right) \Big| \tau_1, \ldots \tau_p,$$

and

$$\left(T_j^0, T_j^1, T_{j'}^0, T_{j'}^1\right) | \tau_1, \dots \tau_p \sim \left(T_j^0, T_j^1, T_{j'}^1, T_{j'}^0\right) | \tau_1, \dots \tau_p$$

The first equality in distribution holds by the same argument for (1) and the fact that swapping $G_{g_k}$ and $\tilde{G}_{g_k}$ does not change $T_{j'}^1$ and $T_{j'}^0$ when $j' \in g_{k'}$, $k \neq k'$. The second equality in distribution can be proved similarly. Then we have

$$\left(T_j^0, T_j^1, T_{j'}^0, T_{j'}^1\right) | \tau_1, \dots \tau_p \sim \left(T_j^1, T_j^0, T_{j'}^1, T_{j'}^0\right) | \tau_1, \dots \tau_p$$

$$\Rightarrow \kappa_j \perp \kappa_{j'} | \tau_1, \dots, \tau_p, \qquad j \in g_k, j' \in g_{k'}, k \neq k'$$

if $y \perp \boldsymbol{G}_{g_k} | \boldsymbol{G}_{-g_k}$ and $Y \perp \boldsymbol{G}_{g_{k'}} | \boldsymbol{G}_{-g_{k'}}$.

In genetic studies, the groups are defined by LD where variants in high correlation reside in the same group. It may be safe to assume that $Y \perp G_j | \boldsymbol{G}_{-g_k} \Leftrightarrow Y \perp \boldsymbol{G}_{g_k} | \boldsymbol{G}_{-g_k}$. Under this additional assumption, the lemma above shows that $\{k_j\}_{j \in R_l, j \in H_0}$ are i.i.d. coin flips conditional on $\{\tau_j\}$. With this lemma, we look at decomposition of FDP as follows:

$$FDP(t_\alpha) = \frac{\#\{j \in H_0 | \kappa_j = 1, \tau_j \geq t_\alpha\}}{1 \vee \#\{j | \kappa_j = 0, \tau_j \geq t_\alpha\}} = \sum_{l=1}^{\phi(t_\alpha)} \frac{\#\{j \in R_l, j \in H_0 | \kappa_j = 0, \tau_j \geq t_\alpha\}}{1 \vee \#\{j | \kappa_j = 0, \tau_j \geq t_\alpha\}}$$

$$= \sum_{l=1}^{L} I\{\phi(t_\alpha) \geq l\} \frac{1 + \#\{j \in R_l | \kappa_j = 1, \tau_j \geq t_\alpha\}}{1 \vee \#\{j | \kappa_j = 0, \tau_j \geq t_\alpha\}} \times \frac{\#\{j \in R_l, j \in H_0 | \kappa_j = 0, \tau_j \geq t_\alpha\}}{1 + \#\{j \in R_l | \kappa_j = 0, \tau_j \geq t_\alpha\}}$$

$$\leq \sum_{l=1}^{L} \frac{I\{\phi(t_\alpha) \geq l\} + \#\{j \in R_l | \kappa_j = 1, \tau_j \geq t_\alpha\}}{1 \vee \#\{j | \kappa_j = 0, \tau_j \geq t_\alpha\}}$$

$$\times \frac{\#\{j \in R_l, j \in H_0 | \kappa_j = 0, \tau_j \geq t_\alpha\}}{1 + \#\{j \in R_l, j \in H_0 | \kappa_j = 1, \tau_j \geq t_\alpha\}}$$

For any fixed $\{v_l\}_{1 \leq l \leq L}$, $\sum_{l=1}^{L} v_l = 1$, if we choose stopping time

$$t_\alpha = \min\left\{t > 0 \left| \frac{I\{\phi(t) \geq l\} + \#\{j \in R_l | \kappa_j = 1, \tau_j \geq t\}}{1 \vee \#\{j | \kappa_j = 0, \tau_j \geq t\}} \leq \frac{\alpha}{1.93} v_l, \ \forall 1 \leq l \leq L \right.\right\}$$

Then we have

$$E\{FDP(t_\alpha)\} \leq \sum_{l=1}^{L} \frac{\alpha}{1.93} v_l E\left\{\frac{\#\{j \in R_l, j \in H_0 | \kappa_j = 0, \tau_j \geq t_\alpha\}}{1 + \#\{j \in R_l, j \in H_0 | \kappa_j = 1, \tau_j \geq t_\alpha\}}\right\}$$

$$\leq \frac{\alpha}{1.93} \sum_{l=1}^{L} v_l E\left\{\sup_t \frac{\#\{j \in R_l, j \in H_0 | \kappa_j = 0, \tau_j \geq t_\alpha\}}{1 + \#\{j \in R_l, j \in H_0 | \kappa_j = 1, \tau_j \geq t_\alpha\}}\right\} \leq \frac{\alpha}{1.93} \sum_{l=1}^{L} v_l \times 1.93$$

$$= \alpha$$

where $\left\{\sup_t \frac{\#\{j \in R_l, j \in H_0 | \kappa_j = 0, \tau_j \geq t_\alpha\}}{1 + \#\{j \in R_l, j \in H_0 | \kappa_j = 1, \tau_j \geq t_\alpha\}}\right\} \leq 1.93$ is by Katsevich and Sabatti (2019) if $\{k_j\}_{j \in R_l, j \in H_0}$ are i.i.d. coin flips conditional on $\{\tau_j\}$.[15] Note that the above is true for any $\{v_l\}_{1 \leq l \leq L}$, $\sum_{l=1}^{L} v_l = 1$. Ideally, we would like to find the optimal $\{v_l\}_{1 \leq l \leq L}$ such that $t_\alpha$ is minimized. While the optimal solution is unknown, we consider two practical solutions:

**Solution 1**. Intuitively, $v_l$ represents the budget that we assign to each row. For rows with potentially more discoveries, we should assign a larger budget. We achieve this by letting

$$v_l = \frac{\sum_{j \in R_l} \tau_j}{\sum_j \tau_j}, \qquad t_\alpha = \min\left\{t > 0 \left| \frac{I\{\phi(t) \geq l\} + \#\{j \in R_l | \kappa_j = 1, \tau_j \geq t\}}{1 \vee \#\{j | \kappa_j = 0, \tau_j \geq t\}} \leq \frac{\alpha}{1.93} v_l, \ \forall 1 \leq l \leq L \right.\right\}$$

Here $v_l$ is a function of $\{\tau_j\}_{1\le j\le p}$. Since the proof is conditioning on the magnitude $\{\tau_j\}_{1\le j\le p}$, $v_l$ is considered as a fixed value.

**Solution 2**. We note that each

$$\frac{I\{\phi(t)\ge l\}+\#\{j\in R_l|\kappa_j=1,\tau_j\ge t\}}{1\vee\#\{j|\kappa_j=0,\tau_j\ge t\}}\le\frac{\alpha}{1.93}v_l,\qquad \forall 1\le l\le L$$

is a sufficient condition, and the FDR control holds for any $\{v_l\}_{1\le l\le L}$, $\sum_{l=1}^{L}v_l=1$. A weaker condition is given by summing up all terms on the left-hand side and right-hand side:

$$\sum_{l=1}^{L}\frac{I\{\phi(t)\ge l\}+\#\{j\in R_l|\kappa_j=1,\tau_j\ge t\}}{1\vee\#\{j|\kappa_j=0,\tau_j\ge t\}}\le\sum_{l=1}^{L}\frac{\alpha}{1.93}v_l$$

which is equivalent to

$$\frac{\phi(t)+\#\{j\in R_l|\kappa_j=1,\tau_j\ge t\}}{1\vee\#\{j|\kappa_j=0,\tau_j\ge t\}}\le\frac{\alpha}{1.93}.$$

The corresponding filter is

$$t_\alpha=\min\left\{t>0|\widehat{FDP}(t)=\frac{\phi(t)+\#\{j\in R_l|\kappa_j=1,\tau_j\ge t\}}{1\vee\#\{j|\kappa_j=0,\tau_j\ge t\}}\le\frac{\alpha}{1.93}\right\}.$$

This stopping time serves as a lower bound for the optimal choice of $\{v_l\}_{1\le l\le L}$, $\sum_{l=1}^{L}v_l=1$. Meanwhile, since $\{v_l\}_{1\le l\le L}$ can be arbitrary as long as $\sum_{l=1}^{L}v_l=1$, this stopping time should not be far from the one given by the optimal $\{v_l\}_{1\le l\le L}$.

Similar to the discussion by Sesia et al. (2020), the correction of the FDR level by the factor 1.93 is required in the proof for technical reasons, although we have observed in numerical simulations that this may be practically unnecessary.[13] At target FDR=0.10, **Solution 1** exhibits FDR=0.042 and **Solution 2** exhibits FDR=0.079 without the 1.93 factors in our genome-wide simulations studies. Based on our extensive simulation studies and real-data applications, we believe that it may be safe to apply the filter even without the 1.93 factor to avoid an unjustified power loss while retaining provable guarantees.
